# Sox2 Regulates Lateral Line Morphogenesis via Yap/Taz-Mediated Mechanotransduction

**DOI:** 10.1101/2025.09.18.677037

**Authors:** Akshai Janardhana Kurup, Aya Mikdache, Nachoi Mokhtar, Patricia Diabangouaya, Gwendoline Gros, Camila Garcia-Baudino, Cristian A. Undurraga, Andres F. Sarrazin, Pedro P. Hernandez

## Abstract

Organ morphogenesis relies on a tightly regulated interplay between cell proliferation, migration, and differentiation. Emerging evidence suggests that mechanical forces act alongside molecular signals to orchestrate tissue patterning, yet how these diverse inputs are integrated remains poorly understood. The zebrafish posterior lateral line offers a powerful *in vivo* model for studying how cellular behaviors and mechanosensitive signaling are spatiotemporally coordinated during organogenesis. Here, we identify the transcription factor Sox2 as a key regulator of lateral line morphogenesis, influencing the positioning, size, and number of neuromasts, the sensory organs of the lateral line. Loss of Sox2 leads to increased lateral line primordium cell proliferation, disrupted ZO1 deposition at the rosettes and smaller neuromasts positioned more posteriorly along the body axis, while Sox2 overexpression produces opposite phenotypes. Sox2 functions in part by repressing Yap/Taz(Wwtr1) signaling in the primordium. Reduced Yap/Taz activity results in more anterior neuromast deposition and premature termination of primordium migration. Furthermore, we show that as the primordium expands through cell-proliferation, increased cell-junction tension activates Yap/Taz, thereby influencing lateral line development. Reducing overproliferation in *sox2^-/-^*embryos diminishes the elevated Yap/Taz activity, supporting a model in which Sox2 limits proliferation to suppress Yap/Taz signaling and ensure proper primordium morphogenesis. These findings show that Sox2, together with the related SoxB1 transcription factor Sox3, regulates lateral line morphogenesis by coordinating primordium proliferation, neuromast size, and Yap/Taz-dependent mechanosensitive signaling.

## Introduction

The coordination between collective cell migration and differentiation is essential for proper tissue morphogenesis during development. These processes are regulated by signaling pathways that act in space and time, but how such signals coordinate migration, proliferation, and differentiation remain poorly understood. In addition to molecular cues, mechanical forces have recently emerged as key regulators of morphogenesis, acting through mechanosensitive pathways that respond to changes in cell density, tension, and tissue architecture. Yet, the integration of mechanical and molecular signals during organ formation is still incompletely characterized (Goodwin and Nelson 2021).

The zebrafish posterior lateral line provides a powerful *in vivo* system to study these questions. The lateral line is a mechanosensory system in fish that detects water movements and vibrations in the environment (Dalle Nogare and Chitnis 2017). Its development occurs externally, making it easily accessible for live imaging and experimental manipulation. The posterior lateral line (pLL) arises from the collective migration of a group of cells, the pLL primordium. This structure migrates along the trunk from the otic vesicle to the tail tip, continuously depositing rosette-shaped clusters of epithelialized cells known as proneuromasts. Within the primordium, leading cells display mesenchymal features, while trailing cells elongate and organize into epithelial rosettes. Proneuromast deposition occurs when trailing cells lose their migratory capacity and are released from the primordium in a stereotyped pattern. Once deposited, proneuromasts eventually differentiate and mature into functional neuromasts, sensory organs composed of centrally located hair cells and surrounding support cells (Metcalfe, Kimmel, and Schabtach 1985, Ghysen and Dambly-Chaudière 2004A). (Ghysen and Dambly-Chaudière 2004B). These hair cells are structurally and functionally analogous to those in the human inner ear (Nicolson 2005).

Previous studies have highlighted the importance of Wnt, Fgf, and Notch signaling in controlling lateral line development. Wnt signaling is active in the mesenchymal leading region and induces Fgf signaling in the trailing domain (Aman and Piotrowski 2008). Wnt signaling supports primordium cell proliferation and maintains the mesenchymal region, whereas Fgf signaling drives rosette formation and neuromast deposition, partly by activating Notch signaling (Gamba et al. 2010; McGraw et al. 2011; Aman, Nguyen, and Piotrowski 2011; Valdivia et al. 2011; Lecaudey et al. 2008; Nechiporuk and Raible 2008; Ernst et al. 2012; Harding and Nechiporuk 2012). Overactivation of Notch in the primordium results in larger neuromasts, independent of increased cell proliferation (Kozlovskaja-Gumbrienė et al. 2017a). The coordinated spatial activity of these pathways is thus critical for the proper development of the lateral line system.

Although the roles of signalling pathways in lateral line development are well characterized, the mechanisms by which distinct transcription factors regulate these pathways remain largely unexplored. Sox2, a neural-specific transcription factor of the SoxB1 family, has been implicated in neuromast hair cell regeneration (Hernández et al. 2007a). Sox2 plays essential roles in early embryonic development, maintenance of neural progenitors, and cell fate specification (Sarkar and Hochedlinger, 2013; Graham et al., 2003). In the zebrafish pLL, Sox2 is expressed in stem-like supporting cells within neuromasts and in the developing primordium (Hernández et al., 2007; Jimenez et al., 2022), yet its role in lateral line development has yet to be elucidated.

Interestingly, studies in other systems, such as mesenchymal stem cells and osteoprogenitors, have shown that Sox2 can directly regulate the expression of the transcriptional co-activators Yap1 (Yes-associated protein 1) (Seo et al. 2013). Also, the interplay between SOX2 and the transcriptional co-activator TAZ (also known as Wwtr1) has been shown to control cancer stemness in multiple tumor contexts (Talukdar et al. 2025; Basu-Roy et al. 2015). Yap1 and Taz are key effectors of the Hippo signaling pathway and central regulators of tissue growth, cell proliferation, and mechanotransduction. When Hippo signaling is active, Yap/Taz are phosphorylated and sequestered in the cytoplasm, keeping them inactive. When Hippo signaling is inactive, Yap/Taz translocates into the nucleus to drive TEAD-dependent transcription. In addition to canonical Hippo inputs, Yap/Taz can also be activated in response to changes in cell shape, cytoskeletal tension, and substrate stiffness (Bossuyt et al. 2013, Yu and Guan 2013, Barry and Camargo 2013, Zhao, Tumaneng, and Guan 2011, Pan 2010, Halder and Johnson 2011). In the lateral line, *yap1* is expressed in the primordium and regulates its size, although *yap1* mutants surprisingly form a normal lateral line without any positioning or neuromast number defects (Agarwala et al. 2015). The mechanisms controlling Yap/Taz activity in this context remain unclear.

Here, we identify Sox2 as a key regulator of lateral line development. Loss of Sox2 function results in increased primordium cell proliferation and size, defects in ZO1 deposition at the rosette center, and a more posterior positioning of neuromasts along the trunk. Similarly, *yap1* overexpression leads to an enlarged primordium and more posteriorly positioned neuromasts. We further demonstrate that *sox2* loss-of-function phenotypes are at least partially mediated by elevated Yap/Taz activity in the primordium. Embryos lacking *yap1* and heterozygous for *taz* (*yap^-/-^;taz^+/-^*) exhibit premature primordium termination, reduced neuromast number, and a more anterior neuromast deposition - partially mimicking the effects of *sox2* overexpression. We also show that proliferation-driven cell-junction tension and actin polymerization promote Yap/Taz activation in the primordium, influencing lateral line development. Finally, reducing overproliferation in *sox2* mutants restores Yap/Taz activity to control levels, supporting a model in which Sox2, together with the related SoxB1 factor Sox3, regulates lateral line morphogenesis by coordinating primordium proliferation, neuromast size, and Yap/Taz-dependent mechanosensitive signaling.

## Results

### Sox2 and Sox3 control lateral line and neuromast patterning

In the lateral line, the Sox2 protein is expressed in mature neuromast stem-like cells that regenerate hair cells following ablation (Hernández et al. 2007b). While *sox2* mRNA expression has been reported in the developing pLL primordium, the specific role of this gene in the patterning and morphogenesis of the pLL remains unexplored. To address its role, we performed a detailed characterization of *sox2*^-/-^. Given that the zebrafish genome encodes *sox3,* a gene with partially redundant functions to *sox2*, we included *sox3^-/-^* and *sox2^-/-^;sox3^-/-^*double mutants in our initial analysis of pLL patterning. To analyze the positioning of neuromasts along the embryonic trunk, we used the *brn3C:mGFP* transgenic line, which labels central neuromast hair cells (Xiao et al. 2005) and *cldnB:lynGFP* transgenic line, which labels all lateral line cells (Haas and Gilmour 2006). At 3 days post-fertilization (dpf), when neuromast deposition by the posterior lateral line (pLL) primordium is complete, *sox2^-/-^* mutants exhibit a more posterior positioning of neuromasts (Fig. 1A). The penetrance of this phenotype was further enhanced in the *sox2^-/-^;sox3^-/-^*double mutants (Fig. 1A). *sox2^-/-^* also showed a trend towards depositing fewer neuromasts compared to controls, although this difference was not statistically significant (Supplementary Fig. 1). Notably, *sox2^-/-^;sox3^-/-^*double mutants showed a significant reduction in the total number of neuromasts in the pLL (Fig. 1A, Supplementary Fig. 1A). In contrast, *sox3*^-/-^ embryos displayed neither defects in neuromast positioning nor changes in neuromast number.

**Figure 1.**
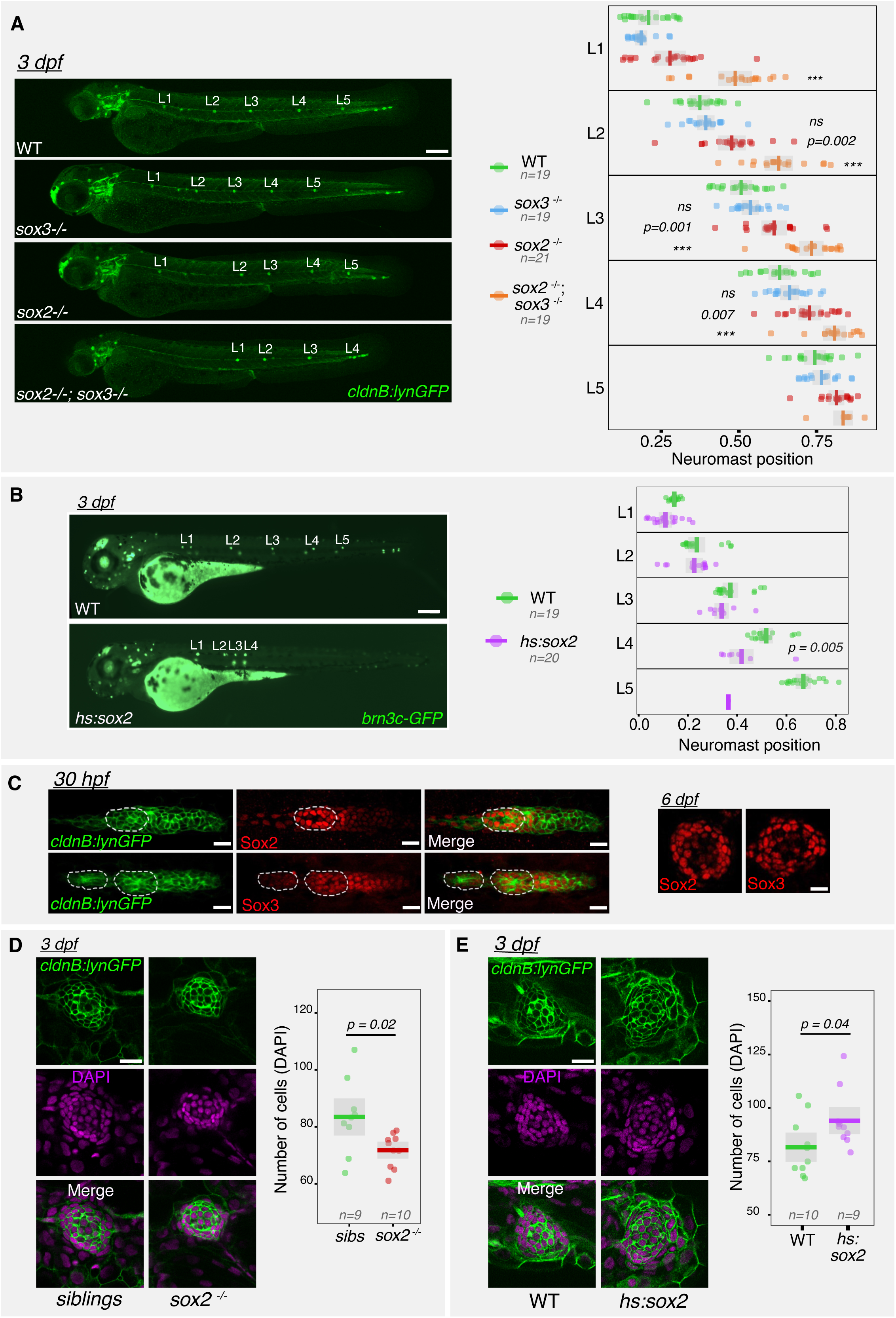
Sox2 controls the number, positioning, and size of neuromasts along the posterior lateral line. **(A)** Quantification of neuromast positions along the embryonic trunk in wild-type, *sox2^-/-^*, and *sox2^-/-^;sox3-/-* embryos. Positions of L1–L5 neuromasts are expressed as a fraction of embryo length. **(B)** Quantification of neuromast positions in *sox2*-overexpressing embryos, expressed as a fraction of embryo length. **(C)** Immunostaining for Sox2 and Sox3 in the posterior lateral line (pLL) primordium and neuromasts. **(D)** Confocal z-stacks of neuromasts in 3 dpf *sox2^-/-^* and sibling embryos stained with *cldnb:lynGFP* and DAPI. Total number of cells per neuromast is quantified. **(E)** Confocal z-stacks of neuromasts in *hsp:sox2* and wild-type embryos stained with *cldnb:lynGFP* and DAPI. Total number of DAPI-positive nuclei per neuromast is quantified. All images are lateral views. In all plots, the mean is shown with standard deviation as error bars. Scale bars: **(A, B)** 200 μm; **(C–E)** 20 μm. Statistical tests were performed using one-way Anova followed by Tukey’s correction in **(A,B)**, Student’s T-test in **(D,E)**; *p*-values are indicated; *ns* = non-significant; \*\*\**P* < 0.001. Data are representative of at least two independent experiments.

To further validate our findings, we took advantage of the heat shock-inducible *hsp-sox2* (Gou et al. 2018a) to analyze the effect of *sox2* overexpression on lateral line development. In *hsp-sox2* heat-shocked embryos, neuromasts were deposited more anteriorly, and fewer neuromasts were deposited on average (Fig. 1B, Supplementary Fig. 1B). We also observed that in these embryos the primordium often failed to reach the tail tip (Fig. 1B)*. sox3* overexpression using the *hsp-sox3* line also resulted in more anteriorly deposited neuromasts, similarly to *sox2* overexpression (Supplementary Fig. 2). These phenotypes inversely mirrored those observed in *sox2* and *sox2;sox3* mutants, supporting a dose-dependent role for these genes in neuromast patterning.

Together, these loss- and gain-of-function analyses identify *sox2* and *sox3* as regulators of neuromast positioning, with partially redundant functions during lateral line development. Since *sox2* single mutants displayed robust defects in neuromast positioning, neuromast size, and primordium development, we used *sox2* mutants as a tractable genetic background for subsequent functional analyses of Sox2/Sox3-dependent lateral line morphogenesis.

Analysis using the *cldnB:lynGFP* transgenic line revealed that neuromasts in *sox2^-/-^* were smaller and contained fewer cells compared to wild-type controls (Fig. 1D). Additionally, upon *sox2* overexpression, individual neuromasts contained more cells (Fig. 1E). Although *sox2^-/-^*did not exhibit a reduction in the number of hair cells within posterior lateral line neuromasts of 3, 4, and 5 dpf embryos, they showed a decrease in the number of support cells (Supplementary Fig. 3). These findings indicate that Sox2 plays a critical role in regulating neuromast size through controlling the support cell number.

### Sox2 controls pLL primordium cell proliferation and proneuromast assembly

Since the migrating pLL primordium gives rise to mature pLL neuromasts, we sought to characterize the morphology of the primordium in *sox2*^-/-^. These mutants exhibited a longer primordium and contained more cells than wild-type embryos (Fig. 2A). Given the elevated cell numbers in *sox2*^-/-^ primordia, we asked whether the primordium cell proliferation rate was also altered. Proliferation within the primordium has been proposed to influence neuromast positioning (Aman, Nguyen, and Piotrowski 2011B; Streichan et al. 2011; Valdivia et al. 2011), although the underlying mechanism remains unclear. EdU incorporation assays revealed a higher ratio of proliferating cells in *sox2*^-/-^ primordia compared to controls (Fig. 2B). Notably, pLL primordium cells in heat-shocked *hsp70:sox2* embryos showed reduced proliferation (Fig. 2C), consistent with a role for *sox2* as a negative regulator of cell division. These findings suggest that Sox2 restricts cell numbers in the primordium by limiting proliferation.

**Figure 2.**
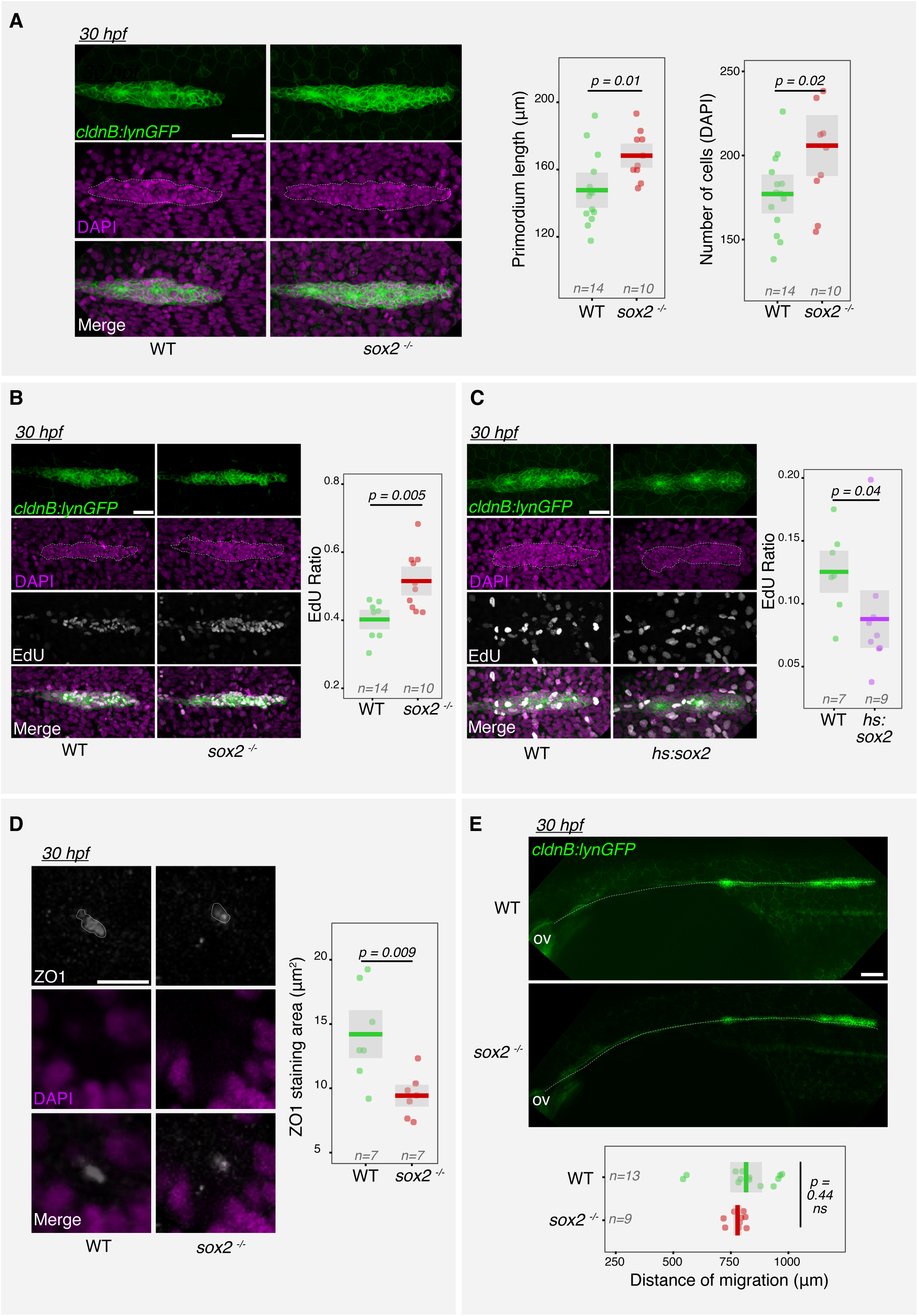
Sox2 controls pLL primordium cell proliferation, length, and patterning. **(A)** Z-projections of pLL primordia in *sox2^-/-^*and WT sibling embryos stained for *cldnb:lynGFP* and DAPI. Total cell number per primordium is quantified. **(B)** Z-projections of EdU-labeled pLL primordia in *sox2^-/-^* and sibling embryos stained with *cldnb:lynGFP* and DAPI. EdU-positive cell ratios are quantified. **(C)** Z-projections of EdU-labeled pLL primordia in *sox2*-overexpressing and control embryos stained with *cldnb:lynGFP* and DAPI. EdU-positive cell ratios are quantified. **(D)** Z-projections of rosettes (R3) in *sox2^-/-^* and sibling embryos immunostained for ZO-1 and DAPI. Quantified regions are indicated with dotted lines. **(E)** Z-projections of *cldnb:lynGFP*-expressing embryos used to measure the distance between the otic vesicle and the primordium tip. All images are lateral views, anterior to the left and dorsal up. In all plots, the mean is shown with standard deviation as error bars. Scale bars: **(A–D)** 50 μm; **(E)** 100 μm. Statistical analyses were performed using the unpaired Student’s t-test in **(A, B, D, E),** Wilcox test in **C**; *p*-values are indicated; *ns* = non-significant, \*\*\**P* < 0.001. Data are representative of at least two independent experiments.

As the pLL primordium undergoes migration and proliferation, cells in its trailing region are assembled into two to three rosettes, referred to as proneuromasts. Cells in the trailing-most rosette (Rosette-3 or R3) lose migratory capacity and are eventually deposited. Rosette assembly influences both the positioning and number of deposited cells. Since *sox2* mutants exhibited mispositioned neuromasts containing fewer cells, we wondered whether rosette assembly was impaired. pLL rosettes are held together by ZO-1, a component of the apically localized tight junction complex (Tornavaca et al. 2015). *sox2* mutants showed smaller ZO-1 foci (Fig. 2D). Since *sox2* mutants showed larger primordia and mispositioned neuromasts, we asked whether this gene also affects primordium migration. At 30 hpf, the distance migrated by the primordium was similar in wild-type and *sox2*^-/-^ embryos, suggesting that this gene does not control migration speed (Fig. 2E). Live imaging revealed a delayed deposition of trailing cells in *sox2* mutants (Video 1), consistent with the analysis shown in Fig. 1A. Overall, our data identify Sox2 as a regulator of primordium cell proliferation and ZO-1 deposition at the rosette center, key processes that influence neuromast positioning.

### Sox2 restricts primordium growth and neuromast patterning by suppressing Yap/Taz activity

Previous studies reported *yap1* expression in the pLL primordium and showed that *yap1* loss reduces primordium cell numbers (Agarwala et al. 2015a; Kozlovskaja-Gumbrienė et al. 2017b). Given that *sox2* mutants exhibited increased numbers of primordium cells, we hypothesized that this phenotype could result from elevated Yap1 activity. To test whether Yap signaling is upregulated in *sox2^-/-^* embryos, we used the *ctgfa:GFP* reporter line (Mateus et al. 2015), which showed strong expression in the pLL primordium (Fig. 3A). We first validated the reliability of *ctgfa:GFP* line in reporting pLL primordium Yap/Taz activity by modulating Yap activity using heat-shock-inducible transgenes: constitutively active Yap *hsp:CA-yap1* increased GFP expression, while dominant-negative Yap *hsp:DN-yap1* reduced it (Supplementary Fig. 4A, B), confirming that *ctgfa:GFP* reports Yap activity in the primordium. In *sox2^-/-^; ctgfa:GFP* embryos, GFP signal was elevated in the primordia (Fig. 3A), indicating an increased Yap activity in the absence of Sox2. To determine whether elevated primordium Yap activity contributes to neuromast mispositioning, we overactivated Yap1 using *hsp:CA-yap1*, which led to a posterior shift in neuromast deposition (Fig. 3B). In contrast, *yap1^-/-^*embryos showed normal neuromast number and positioning (Supplementary Fig. 5), consistent with previous reports (Agarwala et al. 2015a) suggesting that Yap1 alone is dispensable for neuromast positioning. This discrepancy between loss-of-function and overactivation phenotypes led us to consider the possibility of compensatory mechanisms acting in *yap1^-/-^* mutants, as also proposed by a previous study (Agarwala et al. 2015a).

**Figure 3.**
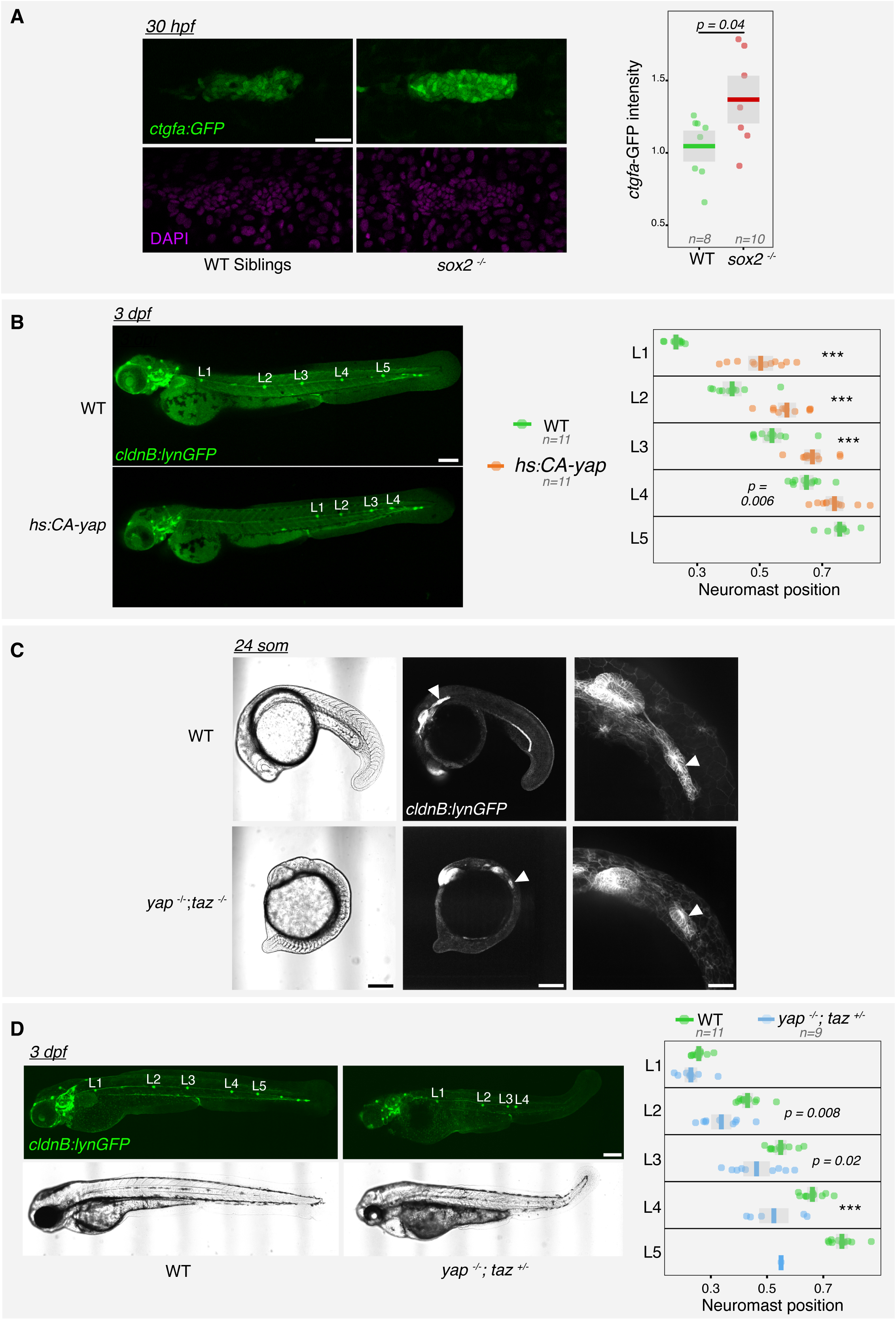
Yap/Taz regulates posterior lateral line morphogenesis. **(A)** Confocal z-stacks of pLL primordia from *ctgfa:GFP* embryos in *sox2^-/-^* and WT sibling backgrounds, stained with DAPI. **(B)** Quantification of neuromast positions in embryos overexpressing a constitutively active form of Yap1. Positions are expressed as a fraction of embryo length. **(C)** Brightfield and fluorescence images of *yap^-/-^;taz^-/-^*embryos showing pLL primordium and posterior body morphology. White arrowheads indicate the pLL primordium. **(D)** Quantification of neuromast positions in *yap^-/-^;taz^+/-^*embryos, expressed as a fraction of embryo length. **(A)** 50 μm; **(B–D)** 200 μm. Statistical analyses were performed using the unpaired Student’s t-test in **(A)** and one-way Anova followed by Tukey’s correction in **(B,D)**; *p*-values are indicated; *ns* = non-significant, \*\*\**P* < 0.001. Data are representative of at least two independent experiments.

Given that Yap1 and Taz can act redundantly in Hippo signaling across vertebrates, including zebrafish (Kimelman et al. 2017), Taz emerged as a potential compensatory factor in the pLL primordium. To investigate the possibility of Taz compensating for Yap1, we examined *yap1^-/-^;taz^-/-^* embryos. At the 24-somite stage, these double mutants displayed posterior body formation defects (Fig. 3C), as previously described (Kimelman et al. 2017), but still formed a pLL primordium, albeit smaller (Fig. 3C), indicating that primordium induction occurs independently of Yap/Taz. However, these embryos were lethal by ∼30 hpf, preventing later analysis of pLL development. We therefore analyzed *yap1^-/-^;taz^+/-^* embryos, which showed anterior neuromast positioning, early termination of the lateral line (Fig. 3D), and fewer neuromasts (Supplementary Fig. 6), resembling the observed *sox2* overexpression phenotype (Fig. 1B). We confirmed by *in situ* hybridization that *cxcl12a* expression is maintained in the horizontal myoseptum of *yap1^-/-^;taz^+/-^*embryos, excluding impaired chemokine-guided primordium migration as a likely cause of the phenotype. (Supplementary Fig. 7). Further, live imaging revealed an early deposition of trailing cells as proneuromasts and disrupted primordium migration in *yap1^-/-^;taz^+/-^* embryos (Video 2). These findings indicate that Yap and Taz act together to regulate neuromast positioning and number.

Since the reduced primordium cell number is known to trigger premature neuromast deposition and early pLL termination (Valdivia et al. 2011), we examined primordium morphology in *yap1^-/-^;taz^+/-^* embryos. These embryos had fewer primordium cells (Fig. 4A) and showed fewer EdU +ve proliferating cells (Fig. 4B). *yap1^-/-^;taz^+/-^* retained normal size of ZO1-positive foci in the trailing-most rosette (R3) (Fig. 4C). We also did not observe increase numbers of apoptotic cells within the primordia of *yap1^-/-^;taz^+/-^* embryos, although cell death appeared elevated among basal epidermal cells in these embryos (Supplementary Fig. 8). Neuromast size was unaffected (Fig. 4D) in the *yap1^-/-^;taz^+/-^*embryos, however, overactivation of Yap1 using *hsp:CA-yap1* resulted in neuromasts containing more cells (Fig. 4E), indicating that regulation of neuromast cell numbers involve suppression of Yap/Taz signaling.

**Figure 4.**
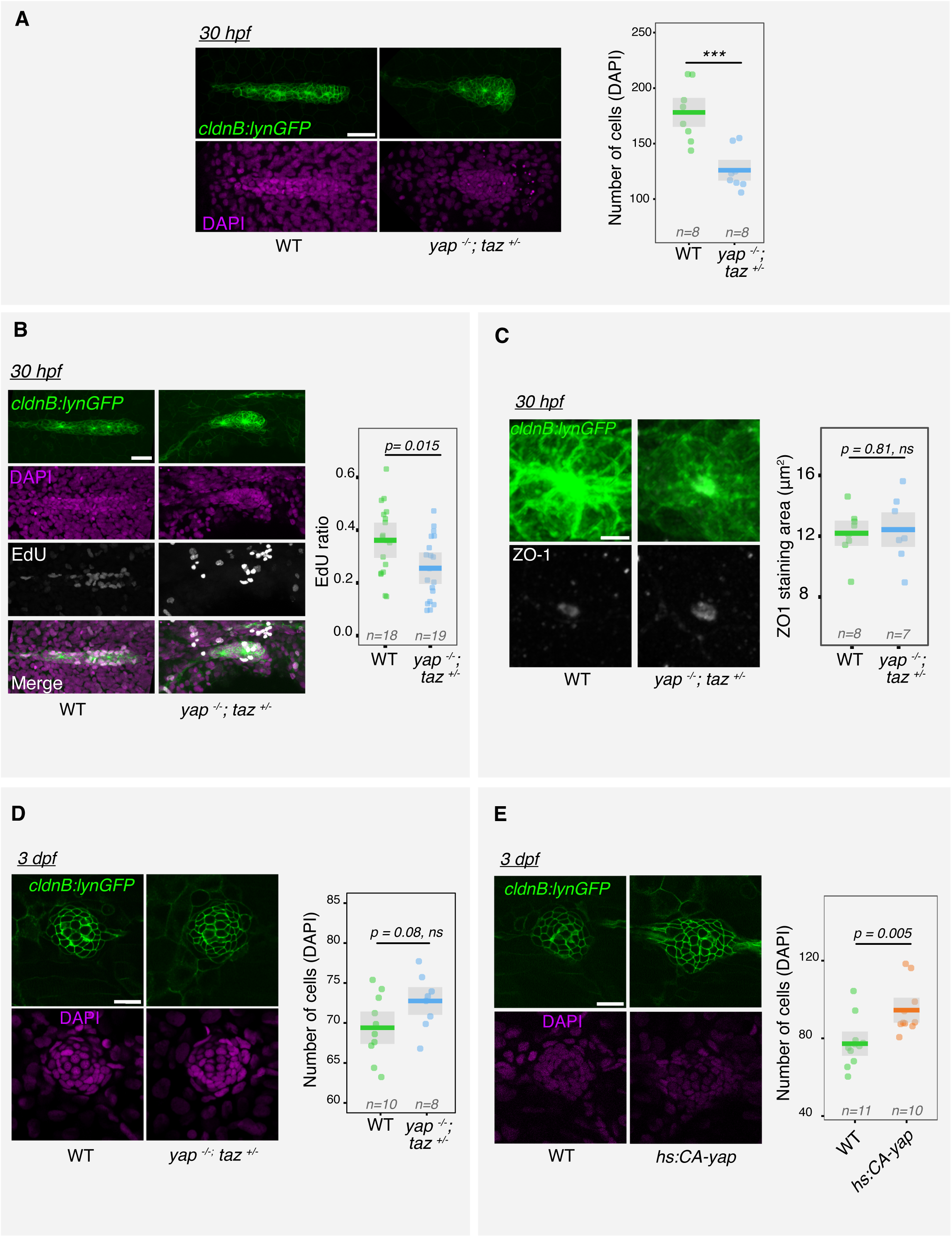
Yap/Taz regulates primordium cell proliferation and neuromast development. **(A)** Z-projections of pLL primordia from *yap^-/-^;taz^+/-^* and control embryos stained with *cldnb:lynGFP* and DAPI. Total number of cells per primordium is quantified. **(B)** Z-projections of EdU-labeled pLL primordia in *yap^-/-^;taz^+/-^* and WT embryos stained with *cldnb:lynGFP* and DAPI. EdU-positive cell ratios are quantified. **(C)** Z-projections of rosettes from *yap^-/-^;taz^+/-^* and sibling embryos stained with ZO-1 and DAPI. All images are lateral views, anterior to the left and dorsal up. **(E)** Confocal z-stacks of neuromasts in 3 dpf *yap^-/-^;taz^+/-^* and sibling embryos stained with DAPI. Total cell numbers per neuromast are quantified. **(F)** Confocal z-stacks of neuromasts in 3 dpf *hsp:CA-yap* and sibling embryos stained with DAPI. Total cell numbers per neuromast are quantified. Scale bars: **(A, B)** 50 μm; **(C, D, E)** 20 μm. Statistical analyses were performed using the unpaired Student’s t-test in (**A, B**)*; p*-values are indicated; *ns* = non-significant, \*\*\**P* < 0.001. Data are representative of at least two independent experiments.

To determine whether the posterior neuromast positioning observed in *sox2* mutants depends on Yap/Taz activity, we first analysed neuromast positioning in *sox2^-/-^; yap1^-/-^* embryos. These embryos displayed a posterior displacement of neuromasts comparable to that observed in *sox2^-/-^* mutants alone, indicating that *sox2^-/-^*phenotype is not solely attributable to *yap1* dysregulation and suggesting that additional factors, potentially *taz*, might also contribute. (Supplementary Fig. 9). We therefore sought to generate *sox2^-/-^; yap1^-/-^; taz^+/-^* embryos by in-crossing *sox2^+/-^; yap1^+/-^; taz^+/-^* adults. However, despite genotyping a large number of adult fish from the corresponding crosses, we failed to recover *sox2^+/-^; yap1^+/-^; taz^+/-^*, suggesting reduced viability or embryonic lethality associated with this genotype. As an alternative approach, we injected a morpholino targeting *sox2* and *sox3* into *yap1^-/-^;taz^+/-^* embryos. The efficiency of this morpholino was validated by its injection resulting in the combined loss of Sox2 and Sox3 expression, as evidenced by the loss of GFP expression in *sox2-p2a-sfGFP* line and Sox3 immunostainings (Supplementary Fig. 10). In a wild-type background, this morpholino produced a statistically significant posterior neuromast shift like the one observed in *sox2^-/-^* and *sox2^-/-^;sox3^-/-^*(Fig. 5A). Importantly, it failed to induce posterior neuromast mispositioning in *yap1^-/-^;taz^+/-^* (Fig. 5A). This indicates that the defects in neuromast positioning observed upon Sox2/3 inhibition requires functional Yap/Taz signaling.

**Figure 5.**
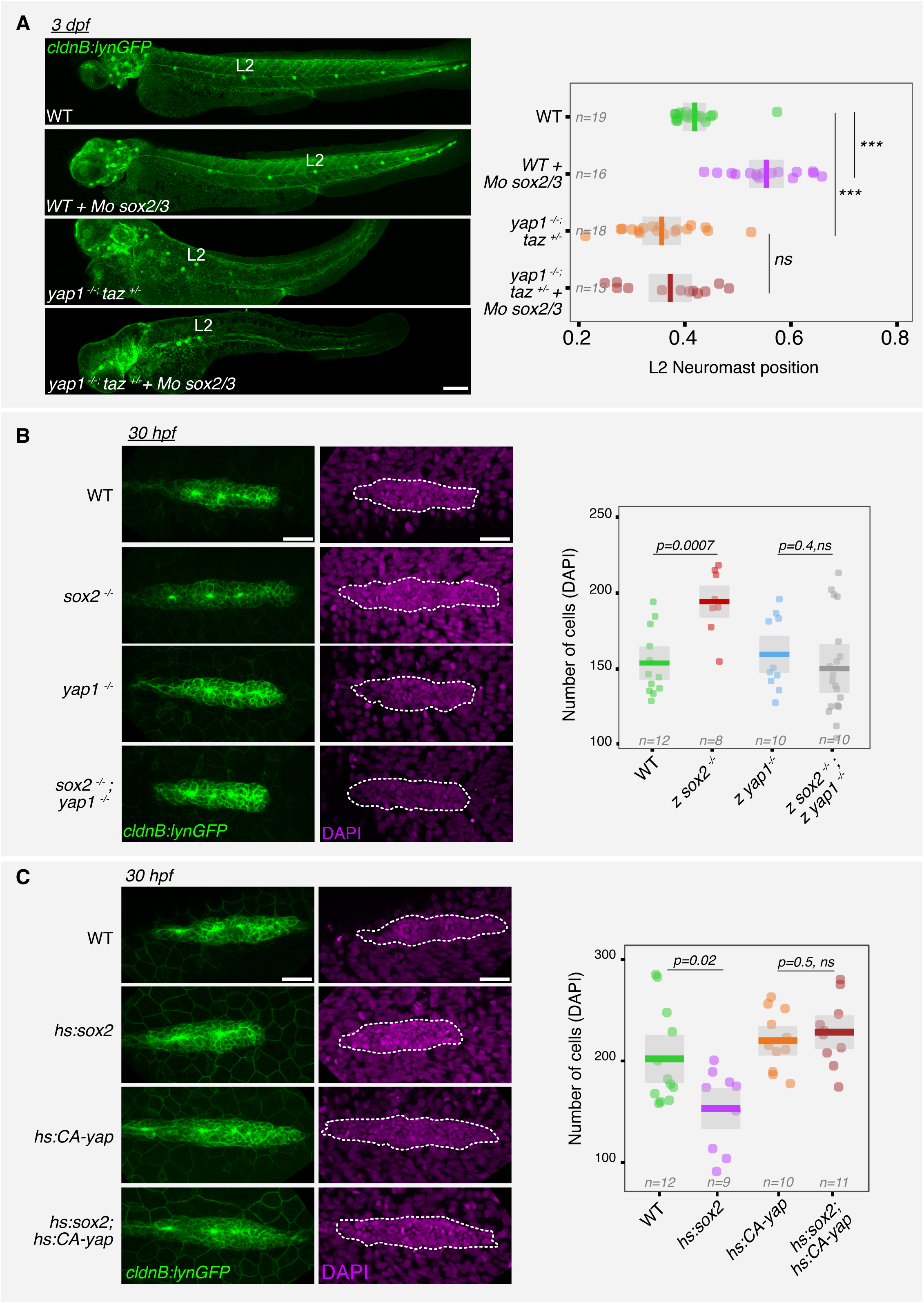
Sox2 suppresses Yap1 activity to regulate neuromast positioning and primordium cell numbers. **(A)** Quantification of neuromast positions along the embryonic trunk in wild-type and *yap1^-/-^; taz^+/-^* embryos injected with MO sox2/3. Positions of L2 neuromasts are expressed as a fraction of embryo length. **(B)** Confocal z-stacks of pLL primordia in *sox2^-/-^*, *yap1^-/-^*, and double mutant embryos stained with DAPI. **(C)** Confocal z-stacks of pLL primordia in WT, *hsp:sox2, hsp:CA-Yap* and *hsp:sox2; hsp:CA-Yap* embryos stained with DAPI and *cldnb:lynGFP*. Scale bars: **A** 200 μm**, (B, C)** 50 μm. Statistical analyses were performed using one-way Anova followed by Tukey’s correction **(A)**, unpaired Student’s t-test in **(B, C)**; *p*-values are indicated; *ns* = non-significant, \*\*\**P* < 0.001. Data are representative of at least two independent experiments.

We next asked whether Sox2 regulates primordium cell numbers through Yap1. The increase in primordium cell numbers in *sox2^-/-^*was no longer observed in *sox2^-/-^;yap1^-/-^* embryos (Fig. 5B), indicating that Sox2 acts upstream of Yap1 to control the number of cells present in the primordium. Consistently, heat-shock-induced *sox2* overexpression reduced primordium cell numbers (Fig. 5C), and this reduction was rescued by co-overexpression of *CA-yap* (Fig. 5C). These results support the idea that Sox2 limits primordium cell numbers, at least in part, by suppressing Yap1 activity. Together, these results identify Sox2 as a negative regulator of Yap/Taz signaling in the primordium and reveal that Yap and Taz cooperatively control neuromast number, positioning, and primordium cell proliferation during lateral line development.

Yap/Taz overactivation did not account for all aspects of the *sox2*^-/-^ phenotype, particularly the reduced number of cells per neuromast, suggesting that Sox2 may act through additional pathways. Given Notch’s known role in neuromast cell number regulation (Kozlovskaja-Gumbrienė et al. 2017a), we investigated its involvement downstream of Sox2. *In-situ* hybridization at 28 hpf showed that while *notch3* expression persisted in the enlarged primordia of *sox2/sox3* morphants, *deltaB* expression was reduced, indicating impaired Notch signaling (Supplementary Fig. 11 A, B). Functionally, overexpression of a dominant-negative *mib1* (*mib1* ΔRF123) reduced neuromast size in wild-type embryos and further exacerbated this phenotype in *sox2^-/-^* embryos (Supplementary Fig. 11 C). Conversely, overexpressing a constitutively activated form of Notch (NICD) was able to restore the size difference between WT and *sox2^-/-^* neuromasts (Supplementary Fig. 11 D). These results suggest that Sox2 promotes Notch signaling to ensure proper neuromast size, acting at least in part through regulation of *deltaB*.

### Proliferation-driven mechanical tension regulates Yap/Taz activity and neuromast positioning

Yap and Taz are well-established mediators of mechanotransduction (Dupont et al. 2011; Wada et al. 2011; Halder, Dupont, and Piccolo 2012; Aragona et al. 2013). For instance, in the embryonic zebrafish heart, myocardial proliferation drives chamber expansion, increasing mechanical tension at endocardial junctions; this tension is sensed by endocardial cells and activates nuclear Yap1 (Bornhorst et al. 2019). Similarly, proliferation-induced compression can trigger Yap activation during mammalian organogenesis, highlighting its sensitivity to mechanical stress (Shroff et al. 2024). In the context of pLL development, it has been shown that inhibiting cell proliferation disrupts neuromast positioning (Aman, Nguyen, and Piotrowski 2011B; Streichan et al. 2011; Valdivia et al. 2011). Given our observation that Yap and Taz influence neuromast-positioning, we hypothesized that proliferative expansion generates mechanical tension, particularly at cell–cell junctions, that activates mechanosensitive Yap/Taz signaling. This activation could serve as a downstream effector linking proliferation-induced mechanical stress to spatial regulation of neuromast deposition.

To characterize the nature of mechanical tension present in cell-cell junctions of the pLL primordium and whether it depends on proliferation, we performed laser ablation of cell junctions in the *cldnb:lynGFP* transgenic line, which labels all primordium cell membranes (Haas and Gilmour 2006). In these embryos, laser ablation of individual junctions caused membranes to recoil and disintegrate, indicating they were under tensile force (Video 3). Quantification of wound width 27 seconds (6 imaging cycles) after ablation confirmed that the cell-cell junctions are under tension (Fig. 6A). In contrast, embryos treated with 40 mM hydroxyurea (HU), a cell cycle inhibitor, showed reduced recoil distances (Fig. 6A), suggesting that cell proliferation contributes to mechanical tension within the primordium. We verified the efficacy of 40 mM HU treatments by performing EdU incorporation and staining, which showed that virtually all primordium cells were EdU+ve, consistent with the reported role for HU in inducing S-phase arrest, during which EdU is incorporated into replicating DNA (Supplementary Fig. 12). To investigate the impact of HU treatment on lateral line development, we treated embryos with 40 mM HU from 20 hpf to 72 hpf. Treated embryos exhibited earlier neuromast deposition compared to controls (Fig. 6B). As Yap/Taz loss of function also leads to premature neuromast positioning, these findings suggest that reduced proliferation might influence Yap/Taz signaling, pointing to a potential mechanistic link between proliferation and Yap/Taz activity. To test whether proliferation contributes to Yap/Taz signaling in the primordium, we treated *ctgfa:GFP* embryos with hydroxyurea (HU). 40 mM HU treatment reduced GFP signal in the primordium (Fig. 6C), indicating that cell proliferation promotes Yap/Taz activity, likely through proliferation-driven mechanical tension.

**Figure 6.**
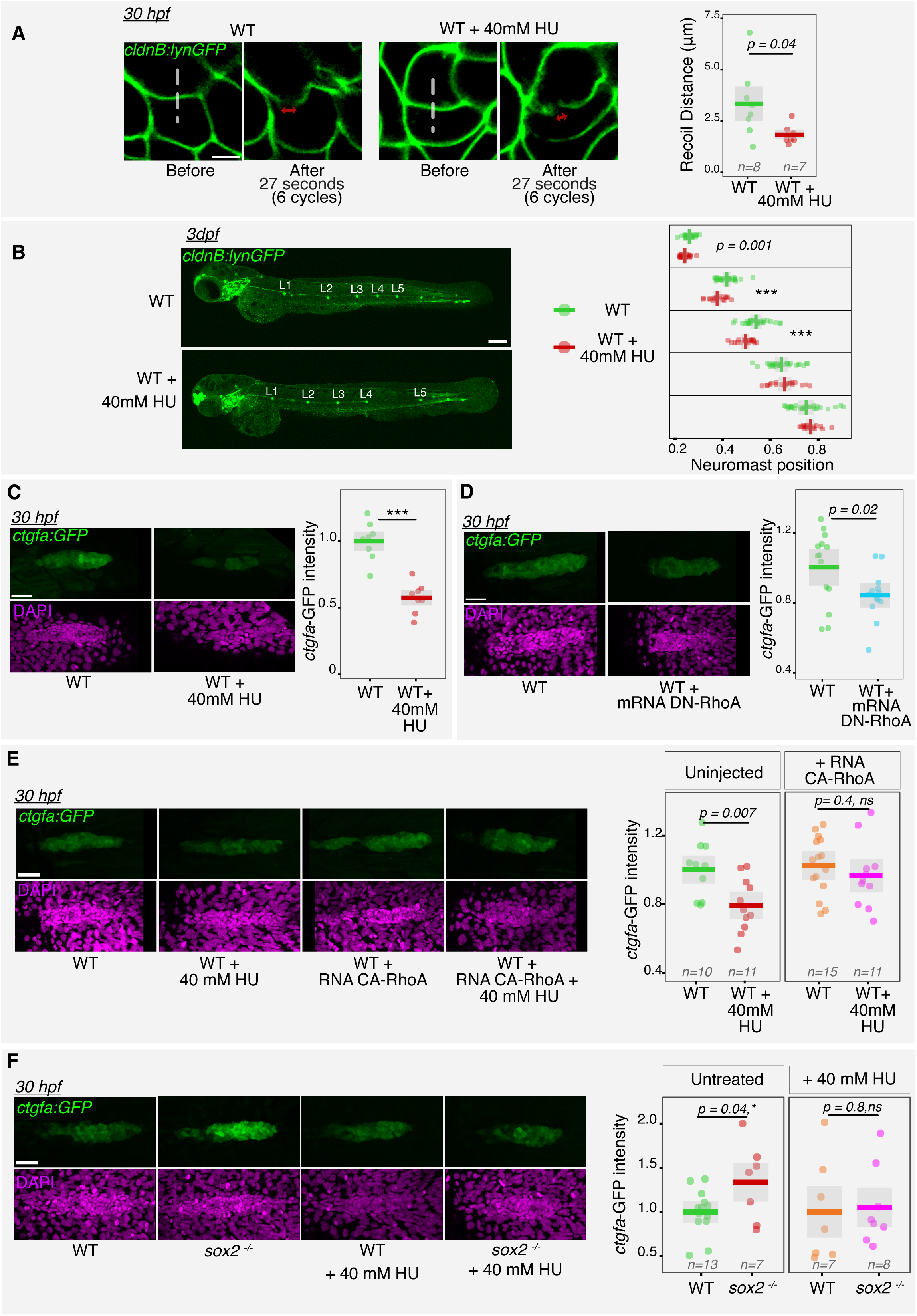
Biomechanical signals regulate Yap/Taz activity in the pLL primordium. **(a)** Images of laser ablation performed on *cldnb:lynGFP* embryos. Dotted lines mark ablation sites; double-headed arrows indicate recoil distances. **(b)** Quantification of neuromast positions in control and hydroxyurea-treated embryos. Positions are expressed as a fraction of embryo length. **(c)** Z-projections of pLL primordia from *ctgfa:GFP* embryos treated with hydroxyurea, stained with DAPI. **(d)** Z-projections of pLL primordia in embryos overexpressing dominant-negative RhoA (DN-RhoA), stained for *ctgfa:GFP* and DAPI. **(e)** Z-projections of pLL primordia in embryos co-treated with hydroxyurea and constitutively active RhoA (CA-RhoA), stained for *ctgfa:GFP* and DAPI. (**f)** Z-projections of pLL primordia in WT and *sox2-/-*embryos co-treated with hydroxyurea stained for *ctgfa:GFP* and DAPI. All images are lateral views, anterior to the left and dorsal up. Scale bars: **(A, C, D, E, F)** 50 μm, **(B)** 200 μm. Statistical analyses were performed using the unpaired Student’s t-test; *p*-values are indicated; *ns* = non-significant, \*\*\**P* < 0.001. Data are representative of at least two independent experiments.

### RhoA acts downstream of proliferation-induced tension to promote Yap/Taz signaling in the primordium

Mechanical stress at cell-cell junctions is known to activate the RhoA/ROCK pathway, promoting F-actin polymerization and downstream YAP/TAZ activation (Elosegui-Artola et al. 2016; Sero and Bakal 2017; Totaro, Panciera, and Piccolo 2018). To test whether RhoA signaling contributes to YAP/TAZ activity in the primordium, we expressed a dominant-negative form of RhoA (DN-RhoA) in *ctgfa:GFP* embryos. DN-RhoA expression led to reduced GFP signal, indicating that RhoA activity is required for YAP/TAZ activation in the pLL primordium (Fig. 6D). Since both inhibition of cell proliferation and suppression of RhoA signaling reduced *ctgfa:GFP* activity, we hypothesized that impaired proliferation limits primordium expansion, thereby reducing junctional tension, RhoA activation, and ultimately Yap/Taz signaling. To test this, we overexpressed a constitutively active form of RhoA (CA-RhoA) in embryos treated with 40mM HU. Unlike wild-type controls, CA-RhoA–expressing embryos did not show a reduction in *ctgfa:GFP* signal upon HU treatment (Fig. 6E), indicating that RhoA functions downstream of proliferation to promote Yap/Taz activity in the primordium.

### Actomyosin contractility is required for Yap/Taz activation in the primordium and enhanced Yap/Taz activity in *sox2* mutants is linked to primordium cell proliferation

Junctional tension can promote actomyosin contractility and increase intracellular tensile forces, thereby contributing to Yap activation (Gaspar and Tapon 2014; Sansores-Garcia et al. 2011). To assess whether proliferation influences cortical F-actin recruitment in the primordium, we used the *act2B:Utrophin-mCherry* transgenic line, which labels filamentous actin at the cell cortex (Behrndt et al. 2012). In 40mM HU-treated embryos, Utrophin-mCherry fluorescence in the primordium was reduced (Supplementary Fig. 13A), indicating diminished F-actin polymerization. To evaluate the role of actomyosin contractility in Yap signaling, we treated *ctgfa:GFP* embryos with 2.5 μM blebbistatin, a myosin II inhibitor. Blebbistatin treatment led to a reduction in GFP expression in the primordium (Supplementary Fig. 13B), indicating that actomyosin-dependent tension drives Yap/Taz activity. Finally, to determine whether the elevated cell proliferation observed in *sox2*^-/-^ contributes to the increased Yap/Taz activity, we treated *sox2^-/-^*; *ctgfa:GFP* embryos with 40 mM HU. This treatment abolished the heightened *ctgfa:GFP* expression seen in untreated *sox2^-/-^* embryos, suggesting that enhanced proliferation drives Yap/Taz overactivation (Fig. 6F). Together, these results support a model in which proliferation-dependent actomyosin contractility promotes Yap/Taz activation in the primordium, and that dysregulated proliferation in sox2 mutants leads to aberrant mechanosensitive signaling.

## Discussion

Morphogenesis requires the precise coordination of cell proliferation, migration, and differentiation, often guided by the integration of molecular signals and mechanical forces. In this study, we use the zebrafish posterior lateral line as a model to dissect how these processes are coupled during sensory organ development. We identify Sox2 as a central regulator of this coordination, acting to ensure proper timing, spacing, and size of neuromasts via its influence on mechanosensitive Yap/Taz signaling.

We characterize Sox2 as a factor controlling the spacing, size of neuromasts, primordium cell proliferation and rosette assembly. Loss of Sox2 disrupts these features, while its overexpression produces complementary, dose-dependent effects, supporting a role for Sox2 in quantitatively controlling primordium behaviour. The enrichment of Sox2 protein in the trailing zone of the primordium suggests spatially regulated activity, although the upstream cues that establish this expression domain remain to be defined. Preliminary results from *sox3* and *sox2;sox3* mutants point to potential redundancy or compensation between these Sox family members, as has been shown in other organs (Gou et al. 2018b).

Furthermore, we identify Sox2 as a negative regulator of Yap/Taz signaling in the primordium. Yap/Taz activity, assessed through a transcriptional reporter, is elevated in *sox2* mutants in a proliferation-dependent manner. Partial loss of Yap/Taz function impairs neuromast deposition and reduces primordium size, and constitutive Yap activation enlarges neuromasts, revealing that Yap/Taz levels must be tightly controlled for proper morphogenesis. These findings expand the known roles of Yap/Taz in the lateral line system, beyond previously reported phenotypes in *yap1* mutants (Agarwala et al. 2015). From previous studies, Sox2 is best known for its roles in maintaining neural and sensory progenitor identity (Sarkar and Hochedlinger, 2013; Graham et al., 2003), but our study expands its function to include control of tissue mechanics via modulation of proliferation and mechanotransduction. This raises the possibility of developmental transcription factors also regulating mechanical inputs and thereby influencing animal development in other contexts. Our data also support a mechanosensitive role for Yap/Taz activation, where proliferation-driven junctional tension within the primordium contributes to Yap/Taz signaling. Inhibition of proliferation reduces junctional tension, cortical actin, and Yap/Taz reporter activity, and blocking actomyosin contractility similarly suppresses Yap/Taz activation. RhoA signaling acts downstream of proliferation and upstream of Yap/Taz in this pathway. Importantly, Yap/Taz overactivation in *sox2* mutants is rescued by proliferation blockade, reinforcing the idea that Sox2 controls Yap/Taz via modulation of biomechanical inputs. Previous work has shown interaction between Sox2 and Yap/Taz in mesenchymal stem cells and osteoprogenitors (Bora-Singhal et al. 2015; Seo et al. 2013). Whether interactions in these model system also involve biomechanical signalling remains an open question.

These findings position the lateral line primordium as a powerful *in vivo* model for studying the interplay between mechanical forces and signaling pathways during organogenesis. While Yap/Taz emerges here as a key mechanosensitive effector, other pathways may also respond to local tissue mechanics. Wnt/β-catenin and Notch signaling, for example, have been shown in other systems to respond to junctional remodeling and proliferation-induced tension (Aoki et al. 2024; Priya et al. 2020). Although a direct interaction between Yap and Wnt effectors has not been established in the lateral line, the similarity between Yap/Taz and *lef1* mutant phenotypes raises the possibility of convergent or sequential regulation. Future studies should investigate how spatial variations in tension within the primordium coordinate multiple signaling outputs to control patterning decisions.

While Sox2 regulates Yap/Taz activity, this alone does not account for all aspects of the *sox2* mutant phenotype. In particular, reduced neuromast size was linked to impaired Notch signaling, as indicated by decreased *deltaB* expression in *sox2/sox3* morphants. Functional experiments confirmed that modulating Notch activity could either exacerbate or rescue the neuromast size defects. These findings suggest that Sox2 coordinates multiple signaling pathways, at least Yap/Taz and Notch, to orchestrate primordium morphogenesis and neuromast positioning.

In this context, a recent reviewed preprint demonstrated that Sox2 interacts with Wnt signaling to regulate neuromast positioning (Palardy et al. 2025). They describe neuromast positioning defects similar to those observed by us and attribute them to dysregulated Wnt signaling. Using expression analyses and pharmacological treatments, the authors demonstrate that Sox2 restricts Wnt activity within the pLL primordium to promote proper neuromast maturation and deposition. In addition, they characterized how multiple primordium signaling pathways regulate Sox2 expression, suggesting that Sox2 functions as an important signaling integrator during lateral line morphogenesis. Our findings extend and complement their observations. In addition to the neuromast positioning defects observed in *sox2^-/-^* embryos, we identify additional roles for Sox2 in regulating neuromast size, support cell number, ZO-1 organization within primordium rosettes, and primordium cell proliferation. Our genetic analyses suggest that several Sox2-associated phenotypes are mediated through Yap/Taz and Notch signaling pathways. Finally, our results raise the possibility that proliferation-associated mechanosensitive changes in sox2 mutants may contribute to Yap/Taz activation during lateral line morphogenesis.

Together, the two studies support a model in which Sox2 acts as a central regulator of posterior lateral line morphogenesis by coordinating Wnt and Yap/Taz signaling. It would be valuable to investigate how Sox2-driven mechanical inputs and Yap/Taz signaling influence Wnt signaling during lateral line development.

Our findings have implications beyond development. Both Sox2 and Yap/Taz are known to contribute to neuromast hair cell regeneration (Hernández et al. 2007a; Ye et al. 2020), yet whether they interact in this context remains unknown. Also, the lateral line provides an accessible model to explore how mechanical cues and transcriptional regulators jointly modulate regenerative competence. Notably, both Sox2 and Yap/Taz are reactivated in tumorigenesis and epithelial repair, where altered tissue mechanics and transcriptional plasticity drive cell proliferation, survival, and migration (Panciera et al. 2020). Investigating their coordinated roles in regeneration may thus uncover conserved principles relevant to cancer biology and tissue repair.

## Materials and methods

### Zebrafish strains and embryo maintenance

All animals used in the study were reared and kept in the zebrafish core facility at the Institut Curie animal facility in accordance with European Union regulations on laboratory animals using protocol numbers: APAFIS#27495-2020100614519712 v14.2.2, #2019_010 and #2022-008 (approved by the French Ministry of Research).

Embryos were raised in E3 medium at 28.5 °C as described by Westerfield (2007). Embryos were staged according to Kimmel et al. 1995 (Kimmel et al., 1995). *sox2* and *sox3* were inactivated using the previously characterized *sox2^x50^* and *sox3^x52^* alleles, respectively (Gou et al. 2018b). For *yap1* and *taz* inactivation, the *yap1^bns19^* and *taz^bns35^* alleles were employed (Astone, M. et al., 2018, Kimelman et al. 2017). The transgenic lines - *Tg(cldnb:lynGFP)* (Haas and Gilmour, 2006), *brn3C:mGFP* (also referred to as *pou4f3:GFP) (Xiao, T et al. 2005), hsp:CA-Yap, hsp:CA-Yap, ctgfa-eGFP (*Mateus R et al., 2015)*, hsp:sox2 (Millimaki, Sweet, and Riley 2010), actb2:utrophin-mCherry* (Behrndt, M. et al., 2012) were previously described. All mutant animals used in the study are zygotic mutants.

Allele-specific PCR was used to identify the WT *sox2* allele (forward primer 5′-AGCGCTTCTTTTCCCAGCAAAG-3′,reverse primer 5′-GCCTCAGCCCAACACCGGGGGCA-3′ as well as the mutant alleles *x50* (forward primer 5′-AGCGCTTCTTTTCCCAGCAAAG-3′, reverse primer 5′-TGTTCCCCGTGCCCCCGGTGTTGAGCC-3′ and *x53* allele of the *sox3* mutants were genotyped as previously described (Gou et al. 2018b). For identifying the *hsp-sox2* transgenic positive animals an allele specific PCR using the primer pair - 5’-GACGAGGTGTTTATTCGCTCT-3’ and 5’-CTTCAGCTCGGTTTCCAT CATG-3’ were used. For genotyping *yap1^Bns19^,* the 160bp region flanking the allele was amplified using the primers *- 5’-CATGTTCGGGGAGACTCCGAG-3’ and 5’-CAGACAAGTAAGAGACCACC-3’* and the presence or absence of the 41Bp *bns19* deletion was detected using the amplicon length. The *taz^bns35^* allele was genotyped as previously described (Kimelman et al. 2017).

For DNA extraction embryos were lysed 20 min at 95 °C in 28.5 µl 50 mM NaOH, and then neutralized by adding 1.5 µl Tris-HCl pH 7.5. PCR amplifications were carried out using GoTaq G2 polymerase (Promega) at 1 mM MgCl2 using the following cycling parameters: 2 min 95 °C - 10 cycles [30 sec 95 °C – 30 sec 65°C to 55°C – 60 sec 72 °C] – 25 cycles [30 sec 95 °C – 30 sec 55 °C – 60 sec 72 °C] – 5 min 72 °C.

### mRNA and Morpholino injections

The sox2/3 Morpholino ( 5’-GCTCGGTTTCCATCATGTTATACAT-3’) (Ogai et al., 2014) was injected at 100 uM. mRNAs were synthesized using the SP6 mMessage mMachine kit (ThermoFisher). mRNAs were injected with 0.2 % phenol red. RNA microinjection was performed using the following constructs and quantities:

Mib1ΔRF123 (16.25 pg) (Zhang et al., 2007b), NICD-pCS2+ (10 pg) (Takke and Campos-Ortega, 1999), CA-RhoA-pCS2+ (0.05pg) (Tahinci E, 2003), DN-RhoA-pCS2+ (25pg) (Tahinci E, 2003).

### Immunostaining

Embryos were fixed overnight at 4 °C in PEM (80 mM Sodium-Pipes, 5 mM EGTA, 1 mM MgCl2) - 4% PFA - 0.04% TritonX100. After washing 2 × 5 min in PEMT (PEM - 0.2% TritonX100), 10 min in PEM - 50 mM NH4Cl, 2 × 5 min in PEMT and blocking in PEMT - 5% FBS, embryos were incubated 2 hrs at room temperature with primary antibodies. Following incubation, embryos were washed in PEMT, blocked in PEMT - 5% FBS, and incubated again with secondary antibodies for 2 hrs. Embryos were again washed in PEMT. The following antibodies were used in the dilutions indicated:

Sox2 - rabbit anti-sox2 (Abcam, ab97959) 1:100

Sox3 - rabbit anti-sox3 (GeneTex, gtx132494) 1:100

Zo1 - Mouse anti-ZO-1 (Invitrogen, ZO1-1A12) 1:250

GFP - Chicken anti-GFP (Aves lab GFP-1020) 1:500

### Microscopy and image analysis

Embryos were mounted in 1% low melting agarose (Sigma) in glass bottom dishes (Mattek) for confocal imaging. The following confocal microscopes were used - Zeiss780, Zeiss880 using a 10X air, 40X water or 40X oil objective. For live imaging, a spinning disc microscope (Nikon) was used. Images were analyzed using Fiji, Napari or custom Python scripts. Quantifications were performed without any knowledge of the sample genotype in a blindfolded way.

For quantifications of neuromast positioning, the horizontal distance of neuromasts - L1-L5 were measured from the head and expressed as a fraction of total embryo length. Quantifications were performed in Fiji.

For cell counting, a Python script based on Stardist 3D was used to segment and detect all the DAPI positive nuclei present in the frame. Following this, an automatic thresholding based approach was used for detecting the pLL primordiums or neuromasts from *cldnB:lynGFP* staining. Finally the Stardist detected nuclei present in the *cldnB:lynGFP* area alone were extracted and counted using the regionprops function of Scikit-image package. Individual segmentations were manually verified and corrected in case of incorrect detections using Napari.

For quantification of fluorescence intensities involving *ctgfa*, using the sum projections of 20 Z-stacks, the primordium area was thresholded and the mean gray values were measured using Fiji. All mean gray values were normalized to the corresponding control group within each experimental replicate.

In experiments involving quantifications of *act2B:Utrophin-mCherry* intensities, The line tool from Fiji was used for measuring the *Utrophin-mCherry* Mean Gray values. Vertical lines from posterior to anterior of pLL primordium were drawn in the region above rosettes. Quantifications were performed on the middle z-stacks.

### Laser membrane dissections and quantifications

A Zeiss880 equipped with a multi-photon Ti::Sapphire laser (Mai Tai HP DeepSee, Spectra Physics). Ablations were performed on 30 hpf embryos using a 40X water objective. A region was selected on a horizontal membrane located at the center of the primordium for ablation. 100% Laser power was used for ablation for 100 cycles, while simultaneous imaging of membrane dynamics was performed using the *cldnB:lynGFP* by exciting using a 488 Laser. The recoil distance was calculated as the distance the membranes have fallen apart following the microdissection in 6 cycles.

### EdU staining and quantification

2nl drops of 500uM EdU solution was injected into the yolk of embryos staged 28 hpf. Injected embryos were incubate at 28.5°C in the dark for 2 hours and fixed in PEM (80 mM Sodium-Pipes, 5 mM EGTA, 1 mM MgCl2) - 4% PFA - 0.04% TritonX100 for 1 hour. Fixed embryos were washed in PEMT three times and blocked in PEMT-5%FBS for 1 hour. After fixation, the embryos were incubated 30 minutes in a staining solution (as per kit protocol) provided as part of the Click-iT™ EdU Cell Proliferation Kit (Invitrogen, #C10337, # C10340, #C10638). Samples were washed and stored in PEMT. EdU stainings were imaged on a Zeiss780 confocal microscope with a 40X objective. EdU ratios were calculated as the fraction of DAPI nuclei in the primordium that are positive for EdU.

### Pharmacological treatments

Hydroxyurea (HU) (Thermo Scientific Chemicals, A10831.03) treatment was performed at the indicated concentrations starting 20hpf. Treatment was perfromed till 30 hfp in experiments involving that stage. For experiments analysing neuromast positioning, the treatment was performed till 72hpf.

Blebbistatin (Thermo Scientific Chemicals, A62237), treatment was performed at a concentration of 2.5 μM starting 20hpf till 30hpf.

### Heat-shocking

Heat-Shocking was performed at 37^0^C for 30 minutes on 20 hpf embryos kept in E3 using a waterbath.

## Acknowledgments

The authors greatly acknowledge the Cell and Tissue Imaging (PICT-IBiSA), Institut Curie, member of the national infrastructure France-BioImaging (https://ror.org/01y7vt929) supported by the French National Research Agency (ANR-24-INBS-0005 FBI BIOGEN). We are grateful to the members of the animal facility of the Institut Curie for zebrafish care. We thank Antonio Jacinto, Bruce Riley, Nicolas David, Erez Raz, Phong Nguyen, Jean-Leon Maitre, SIlvia Fre, Yun-Jin Jiang, and Naoki Mochizuki for sharing of fish lines and reagents. Also, many thanks to Miguel Allende from the Universidad de Chile for this initial contribution to this project. We sincerely thank Yohanns Bellaïche for his thoughtful comments on the manuscript.

## Funding

This work was supported by the Institut Curie, INSERM, and CNRS, and the grants listed below.

Laboratoire d’Excellence (Labex) DEEP (ANR-11- LBX-0044, ANR-10-IDEX- 0001-02 PSL) (PPH)

Ville de Paris Emergence Program (2020 DAE 78) (PPH)

FRM amorçage (AJE201905008718) (PPH)

ATIP-Avenir Starting Grant R21045DS (PPH)

ERC-StG Cytok-Gut 101041422 (PPH, SR, AJK, YS)

## Author contributions

Conceptualization: PPH, AJK

Methodology and Investigation: AJK, AM, CAU, NM, and AFS designed and conducted experiments

Visualization: AJK, AM, CAU

Funding acquisition: PPH

Writing original draft: AJK, PPH

Writing review & editing: AJK, PPH

## Data and materials availability

The new reagents and datasets generated during the current study are available from the corresponding author upon request.

## Supplementary Materials

**Supplementary Figure 1.**
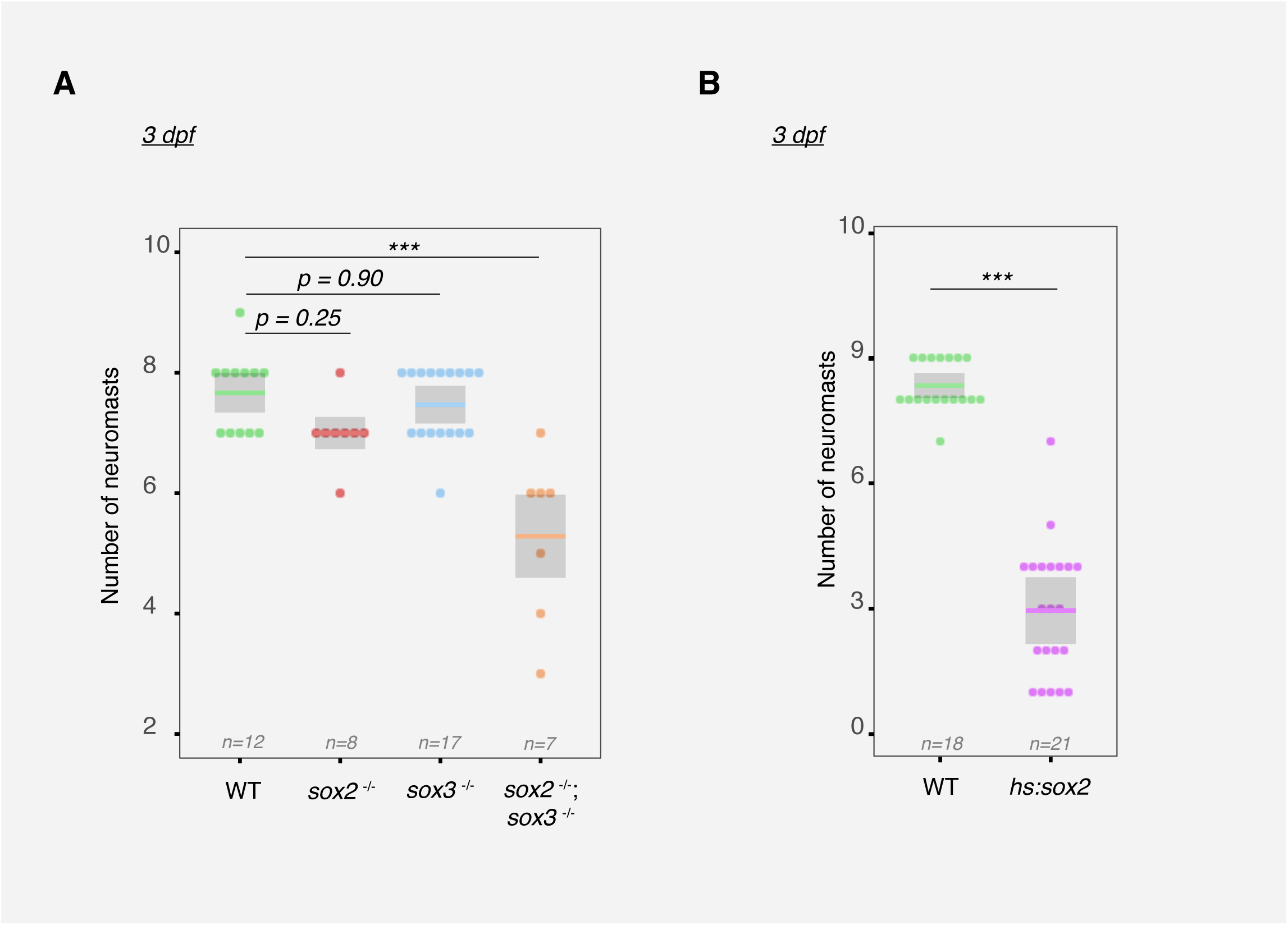
Sox2 and Sox3 control the total neuromast deposited. **(A)** The total number of neuromasts deposited in WT, *sox2^-/-^, sox3^-/-^ and sox2^-/-^;sox3^-/-^*embryos. Part of the embryos shown in main Fig 1a were used for quantification. **(B)** Total number of neuromasts deposited in WT and *hs:sox2* embryos. Embryos shown in Fig 1b were used for quantification. Terminal neuromasts were also counted for quantifications. Statistical analyses were performed using one-way Anova followed by Tukey’s correction in (A) and unpaired Student’s t-test in (B) and p-values are indicated; *ns* = non-significant, ***P < 0.001. Data are representative of at least two independent experiments.

**Supplementary Figure 2.**
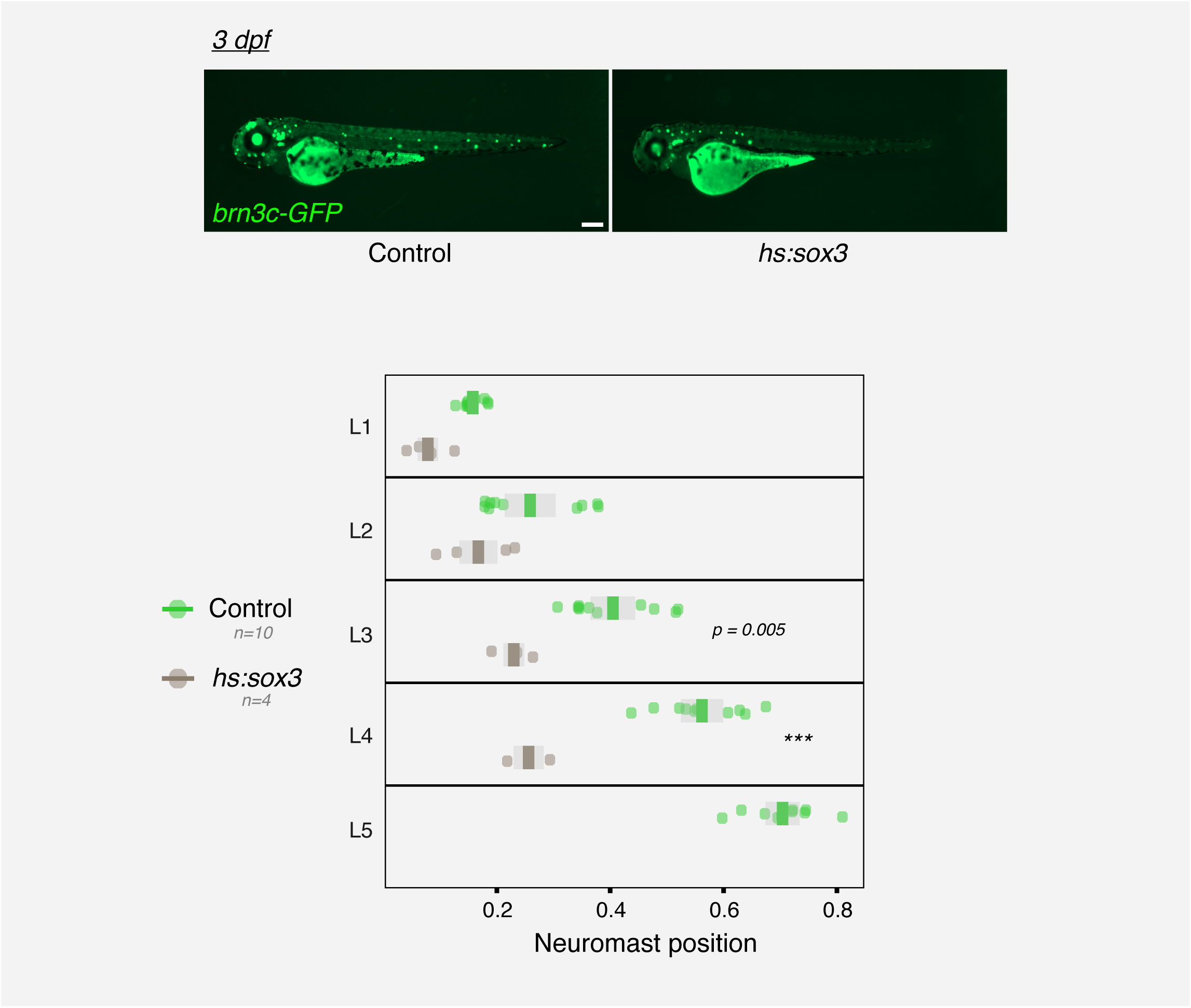
Sox3 overexpression results in more anterior neuromast positioning. Quantification of neuromast positions along the embryonic trunk in Control and *hs:sox3*. Positions of L1–L5 neuromasts are expressed as a fraction of embryo length. Scale bars: 200 μm. Statistical analysis was performed using one-way Anova followed by Tukey’s correction. *ns* = non-significant; ***P < 0.00.1. Data are representative of at least two independent experiments.

**Supplementary Figure 3.**
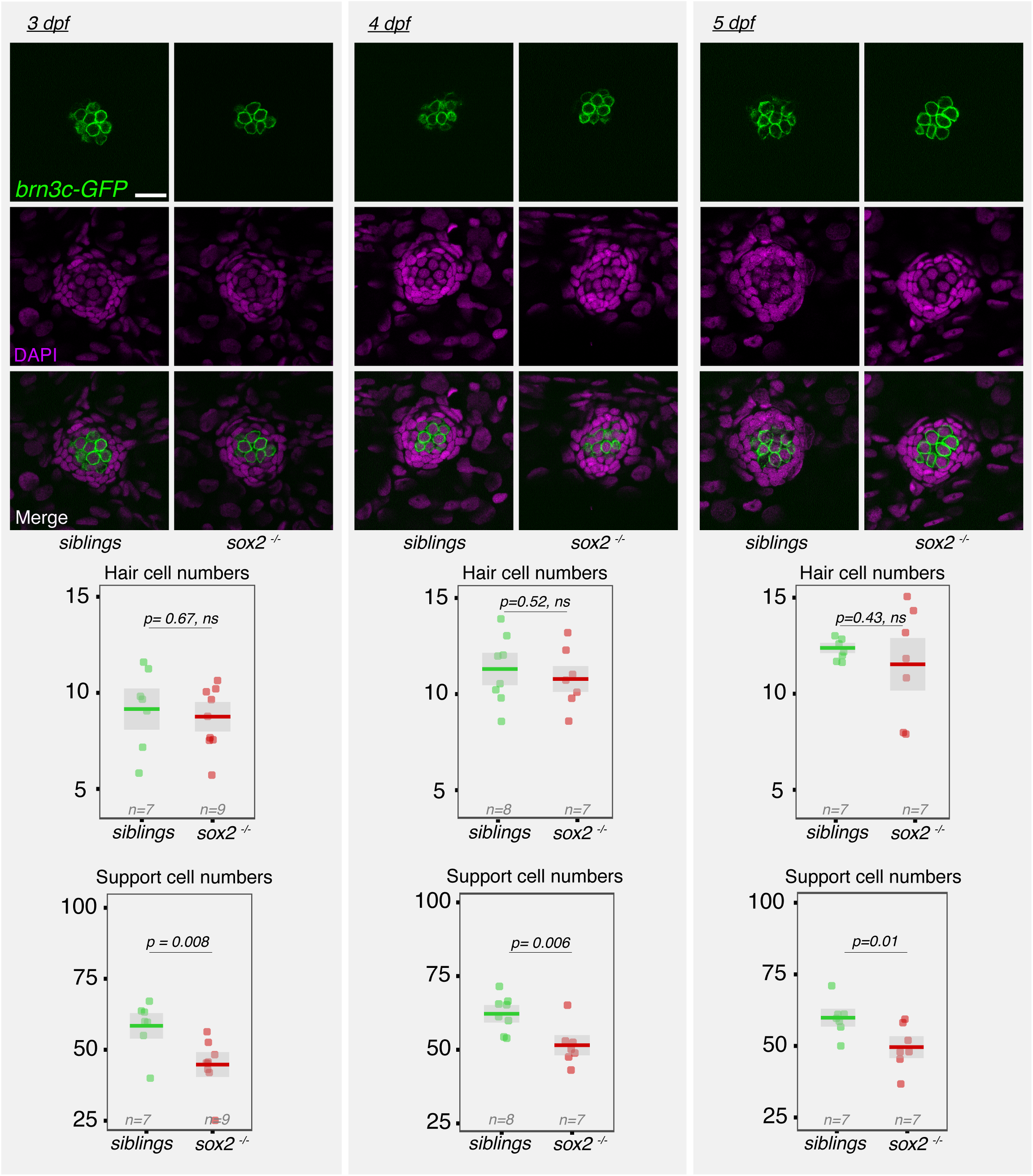
Sox2 regulates the number of neuromast support cells but not hair cells. Confocal z-stacks of neuromasts in 3, 4 and 5 dpf *sox2^-/-^*and sibling embryos expressing *brn3c-GFP* stained with DAPI. *brn3c-GFP* expressing cells were counted as hair-cells. Support cells were counted based on DAPI based nuclei-morphology. Scale bars: 20 μm. Statistical analyses were performed using the unpaired Student’s t-test and p-values are indicated, *ns* = non-significant. Data are representative of at least two independent experiments.

**Supplementary Figure 4.**
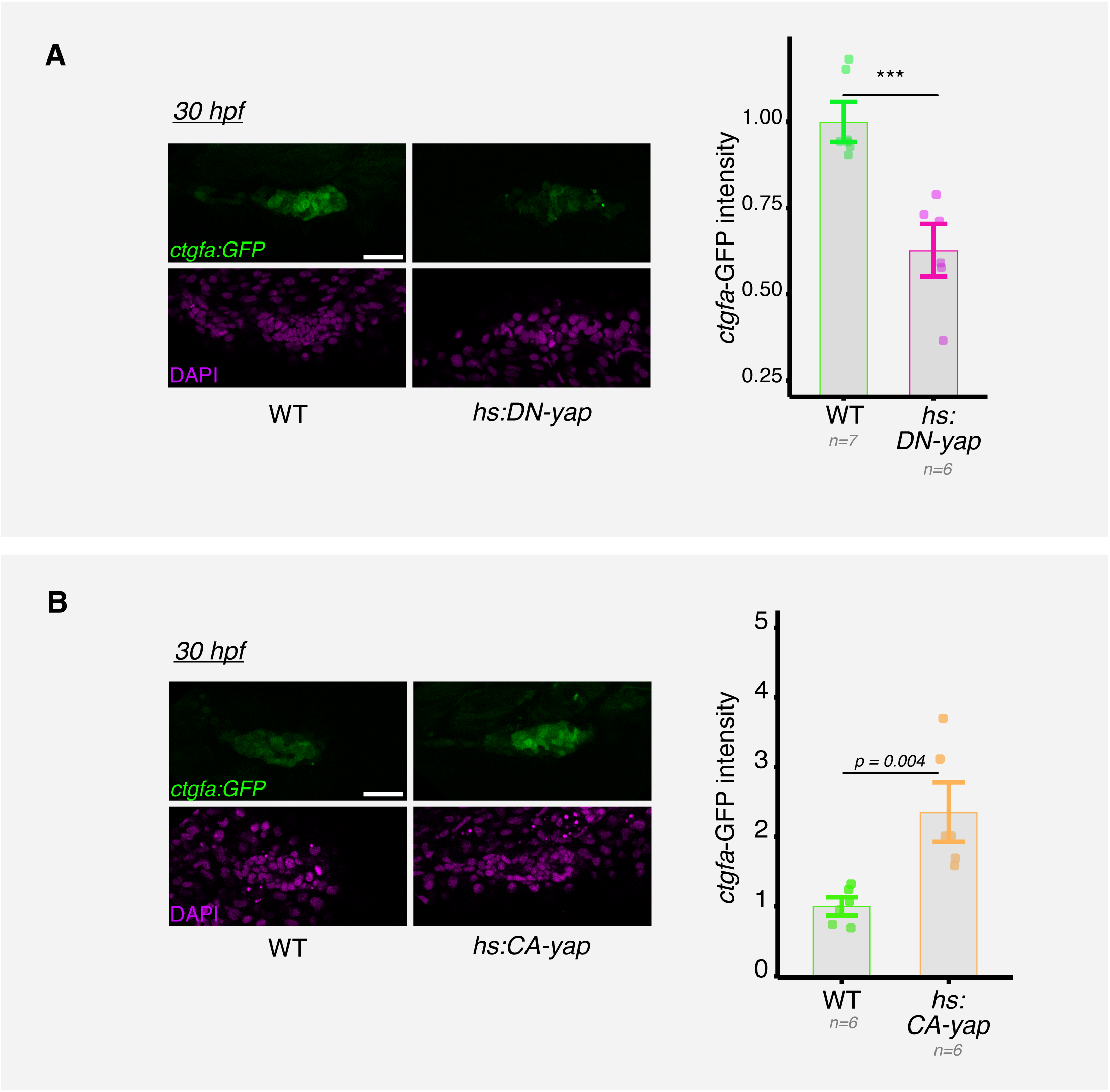
Primordium *ctgfa:GFP* expression responds to Yap activity. **(A)** Confocal z-stacks of pLL primordia from *ctgfa:GFP* embryos in WT and *hs:DN-yap* conditions, stained with DAPI. **(B)** *ctgfa:GFP* expressing primordia in WT and *hs:CA-yap.* Sum intensities from 20 z-stacks were used for quantification. All images are lateral views, anterior to the left and dorsal up. Scale bars: (A, B) 50 μm. Statistical analysis was performed using the unpaired Student’s t-test and the p-values are indicated; ***P < 0.001. Data are representative of at least two independent experiments.

**Supplementary Figure 5.**
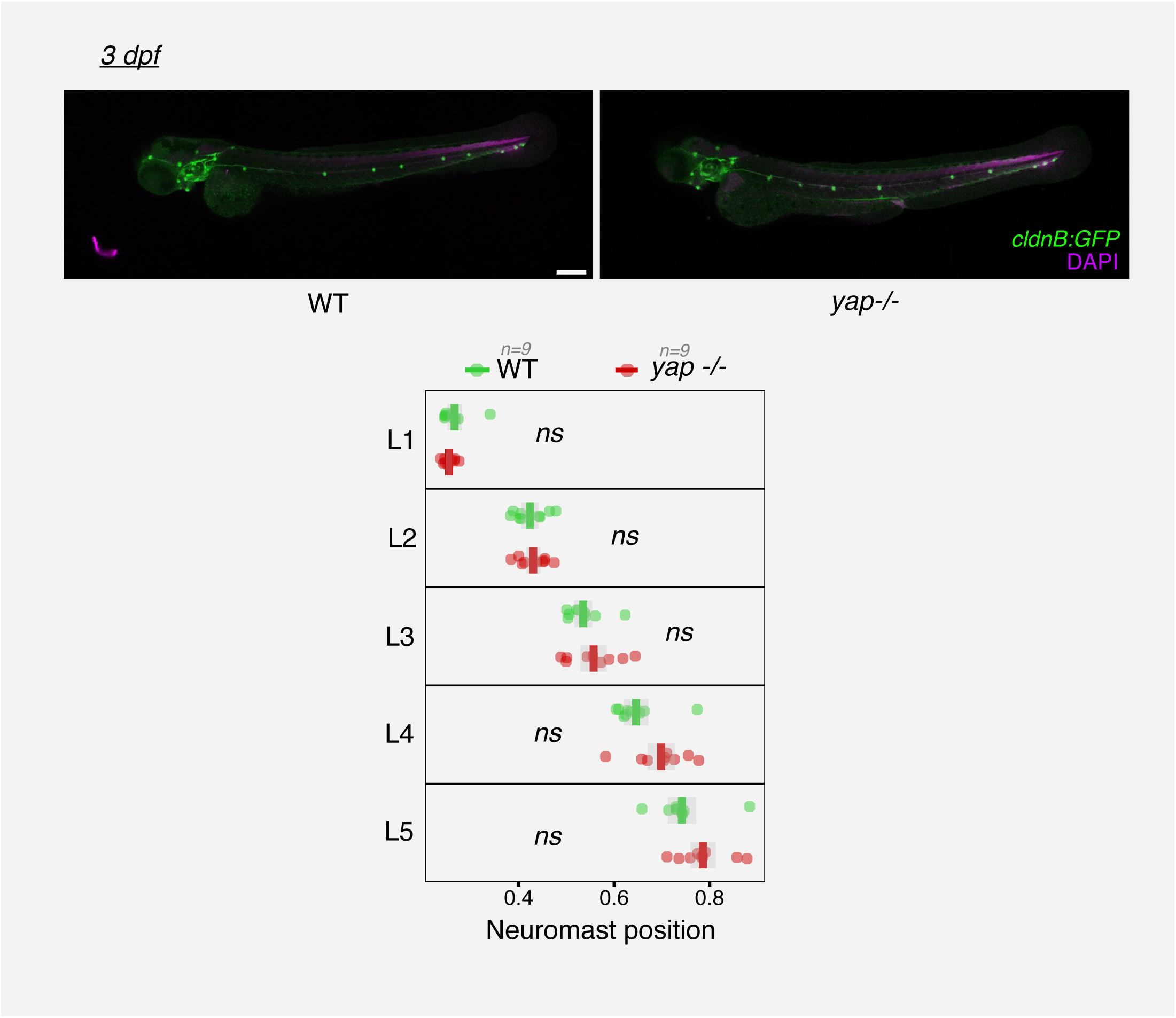
Neuromast positioning is unaffected in *yap1^-/-^*embryos. This experiment is part of the dataset presented in Supplementary Figure 9 and is shown here for illustrative purposes. Quantification of neuromast positions along the embryonic trunk in wild-type and *yap1^-/-^*. Positions of L1–L5 neuromasts are expressed as a fraction of embryo length. Scale bars: 200 μm. Statistical analysis was performed using one-way Anova followed by Tukey’s correction; *ns* = non-significant. Data are representative of at least two independent experiments.

**Supplementary Figure 6.**
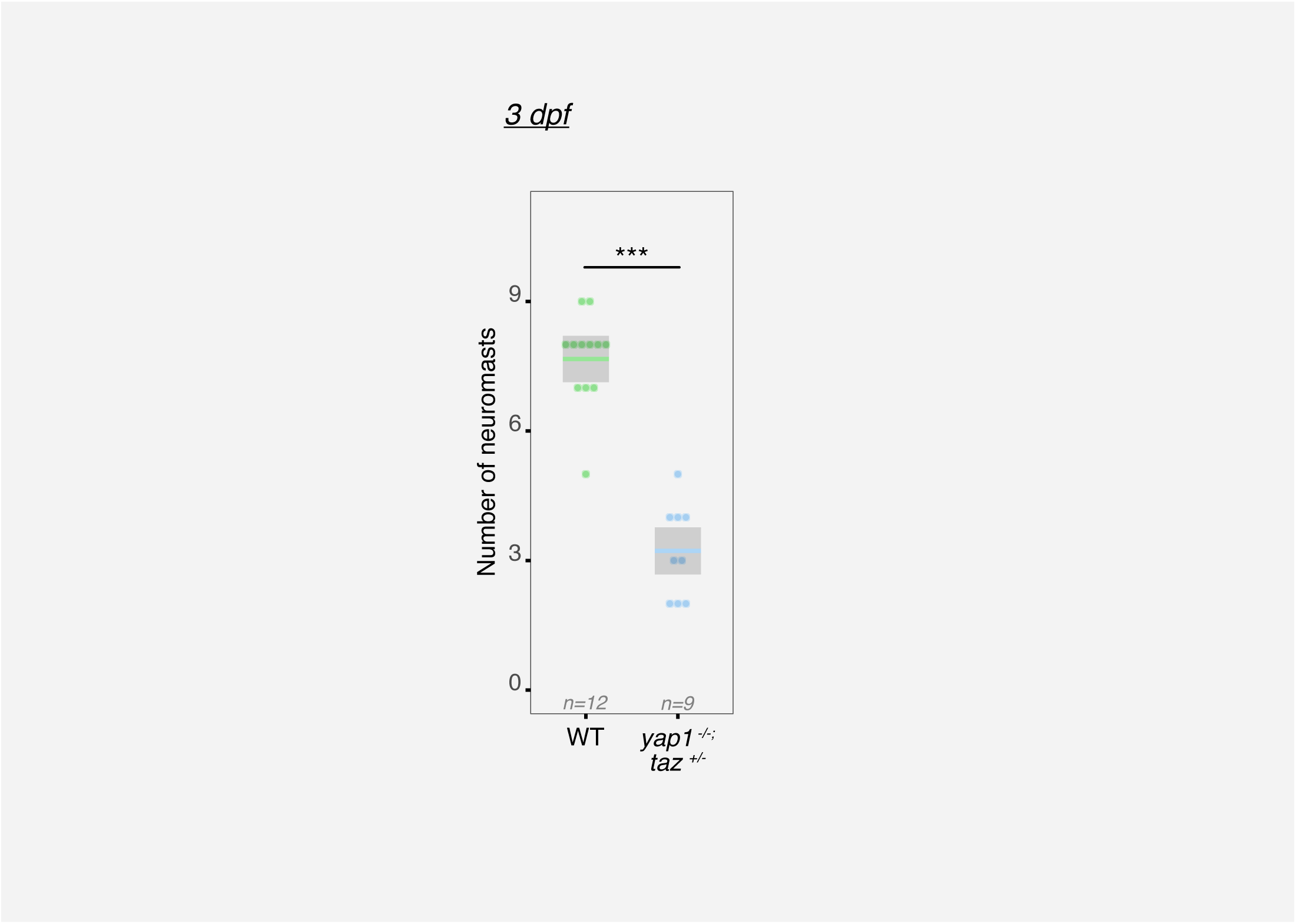
Yap/Taz signaling controls the total number of neuromasts deposited. **(a)** The total number of neuromasts deposited in WT and *yap1^-/-^; taz^+/-^* embryos. WT and *yap1^-/-^; taz^+/-^* embryos shown in main Fig 3d were used for quantification. Terminal neuromasts were also counted for quantifications. Statistical analysis was performed using the unpaired Student’s t-test; ***P < 0.001. Data are representative of at least two independent experiments.

**Supplementary Figure 7.**
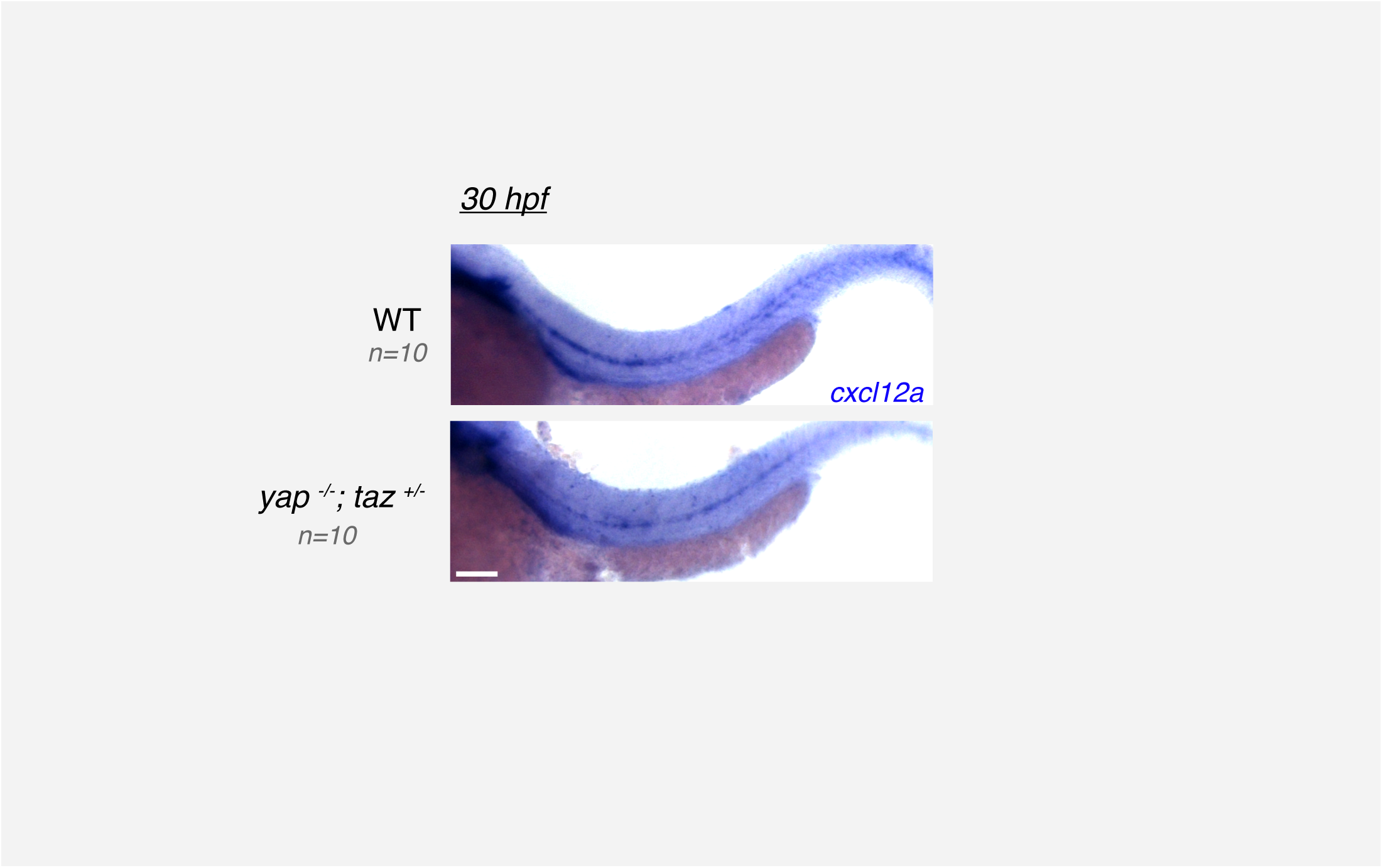
*cxcl12a* expression is unaffected in *yap1^-/-^; taz^+/-^* embryos. *cxcl12a* in situ hybridization in 30hpf WT and *yap1^-/-^; taz^+/-^* embryos. Scale bars: 200 μm. Data are representative of at least two independent experiments.

**Supplementary Figure 8.**
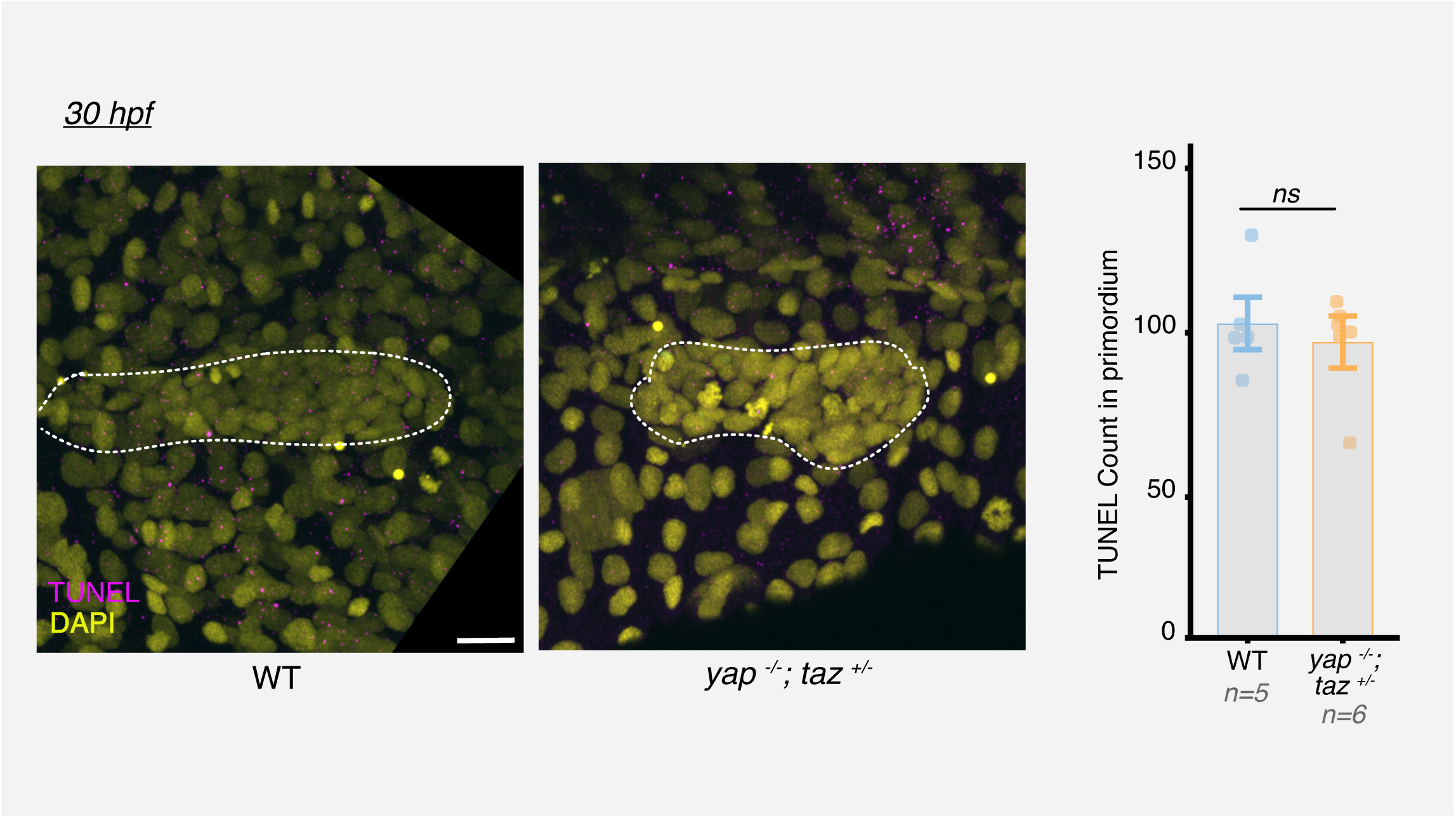
*yap1^-/-^; taz^+/-^* embryos do not show aberrant lateral line primordium cell death. Maximum intensity projections of WT and *yap1^-/-^; taz^+/-^* embryos stained with TUNEL. Total number of TUNEL fragments in the entire primordium was used for quantifications. Scale bars: 20 μm. Statistical analysis was performed using the unpaired Student’s t-test; *ns* = non-significant. Data are representative of at least two independent experiments.

**Supplementary Figure 9.**
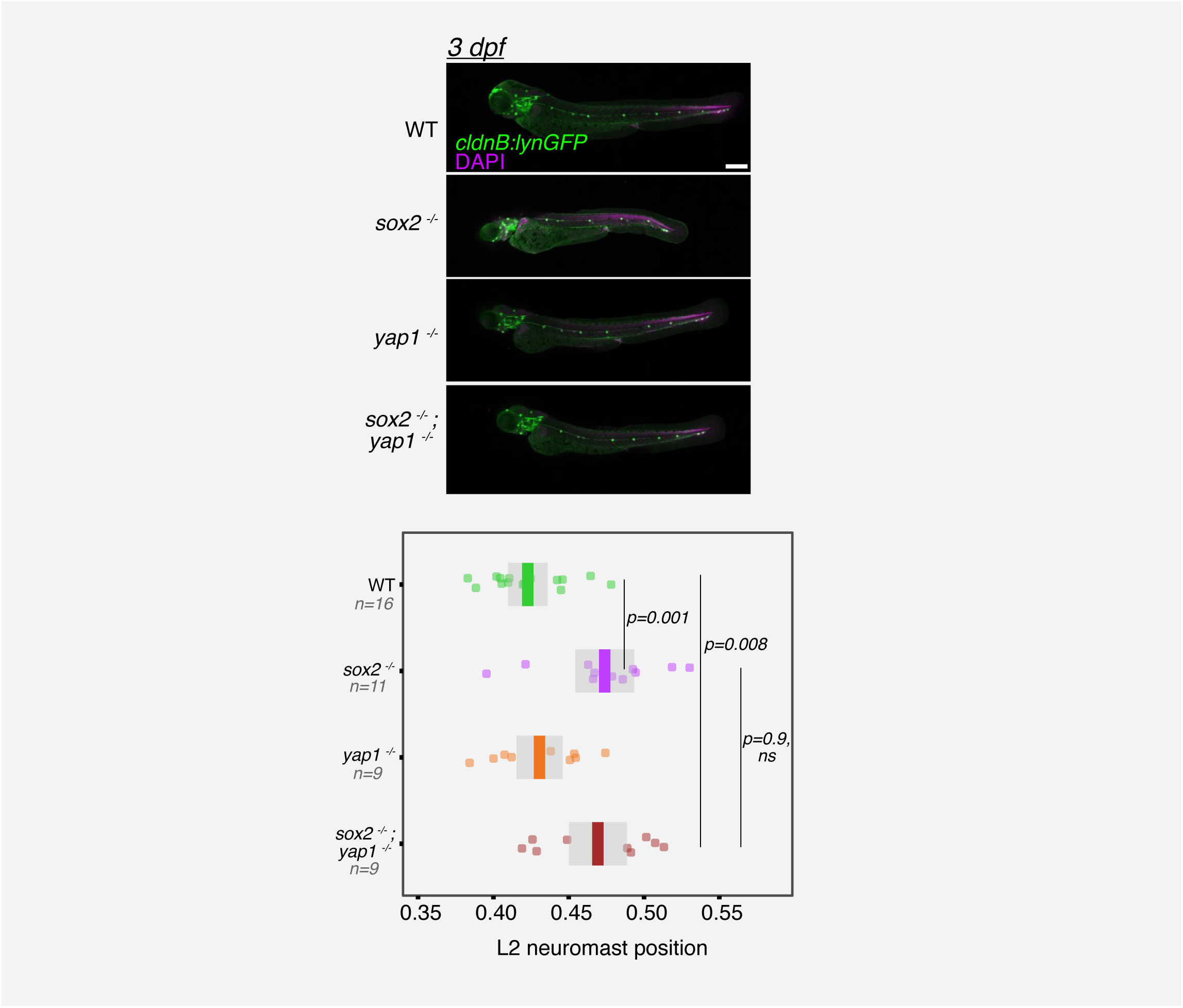
Defective neuromast positioning in *sox2^-/-^* is not solely dependent on *yap1* activity. Part of this dataset has been presented in Supplementary Figure 5 for illustrative purposes. Quantification of neuromast positions along the embryonic trunk in wild-type, *sox2^-/-^*, *yap1^-/-^, sox2^-/-^; yap1^-/-^* embryos. Positions of L1–L5 neuromasts are expressed as a fraction of embryo length. Scale bars: 200 μm. Statistical analyses were performed using one-way Anova followed by Tukey’s correction and p-values are indicated; *ns* = non-significant. Data are representative of at least two independent experiments.

**Supplementary Figure 10.**
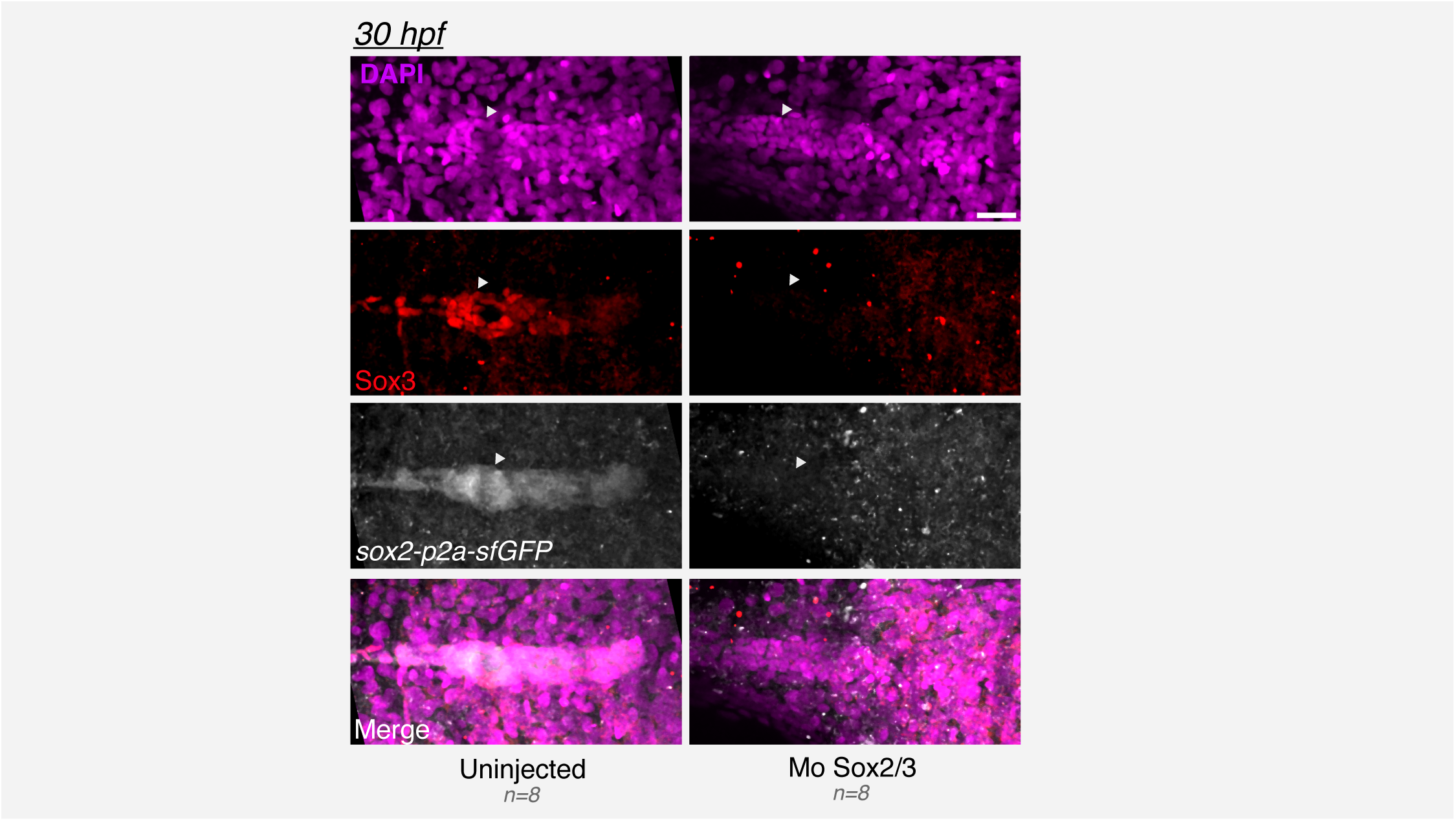
Sox2/3 morpholino injection results in decreased Sox2 and Sox3 protein levels. Maximum-intensity projections of the lateral line primordium in *sox2-p2a-sfGFP* embryos immunostained for Sox3. control and Sox2/3 morpholino-injected conditions shown. Arrowhead points to the trailing region of the primordium. Scale bars: 20 μm. Data are representative of at least two independent experiments.

**Supplementary Figure 11.**
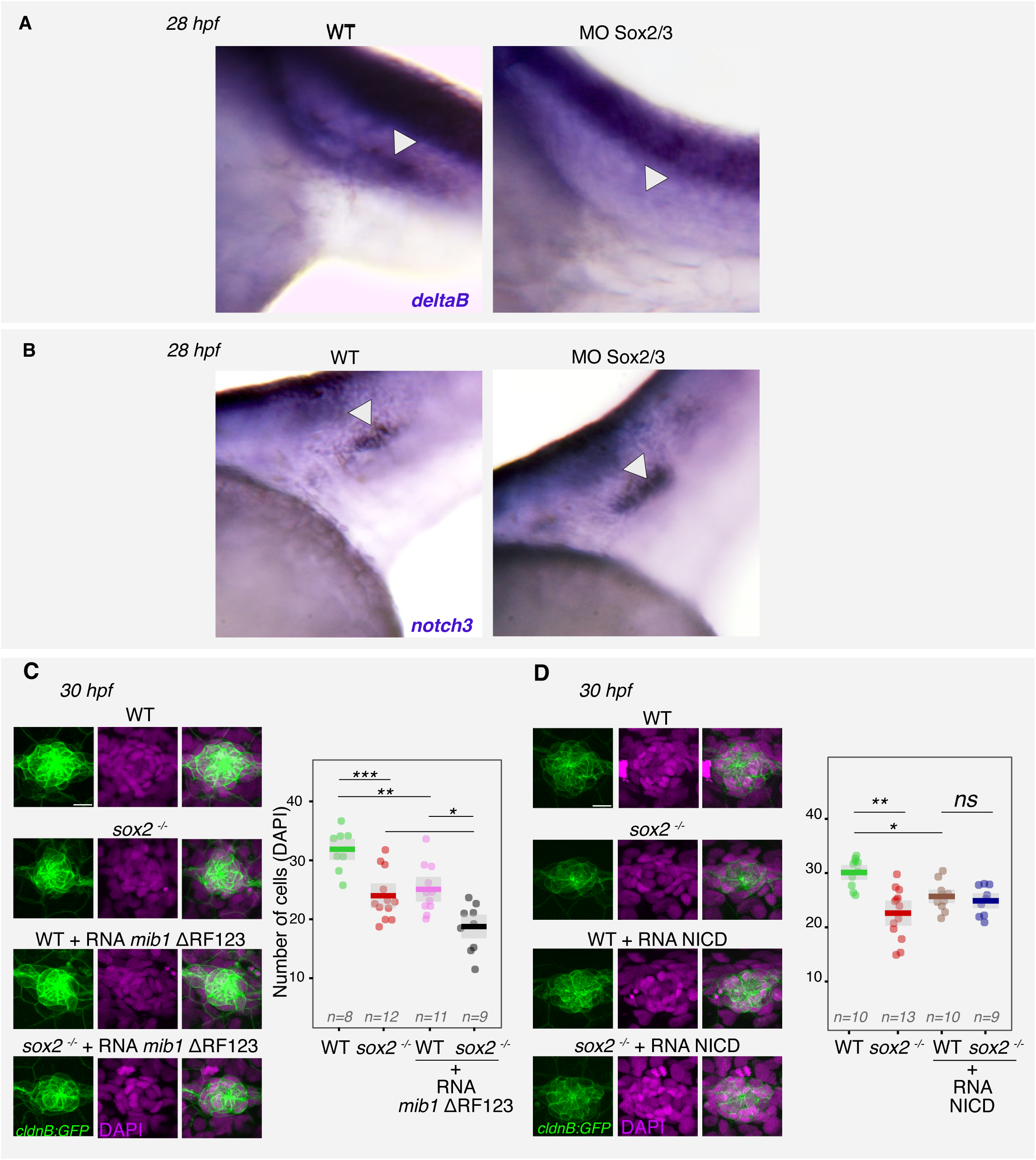
Sox2 may regulate neuromast size by modulating Delta-Notch signaling. (A) Lateral views of *deltaB* expression at 28hpf in WT and Mo Sox2/3 injected emrbryos, arrowheads pointing to the primordium. (B) Lateral views of *notch3* expressing in WT and Mo Sox2/3 morphant embryos at 28hpf, arrowheads pointing to the primordium. (C) Z-projections of L1 neuromasts stained for *cldnB:GFP* and DAPI in the indicated conditions. Images are lateral views, dorsal up. (D) Z-projections of L1 neuromasts stained for *cldnB:GFP* and DAPI in the indicated conditions. Images are lateral views, dorsal up. For quantification, total DAPI cells in the neuromasts were counted. Scale bars: (C and D) 10 μm. Data are representative of at least two independent experiments. Statistical analysis was performed using one-way Anova followed by Tukey’s correction in (c) and (d). All images are lateral views. *ns* = non-significant; *P < 0.05, **P < 0.01, ***P < 0.001, and ****P < 0.0001. Data are representative of at least two independent experiments.

**Supplementary Figure 12.**
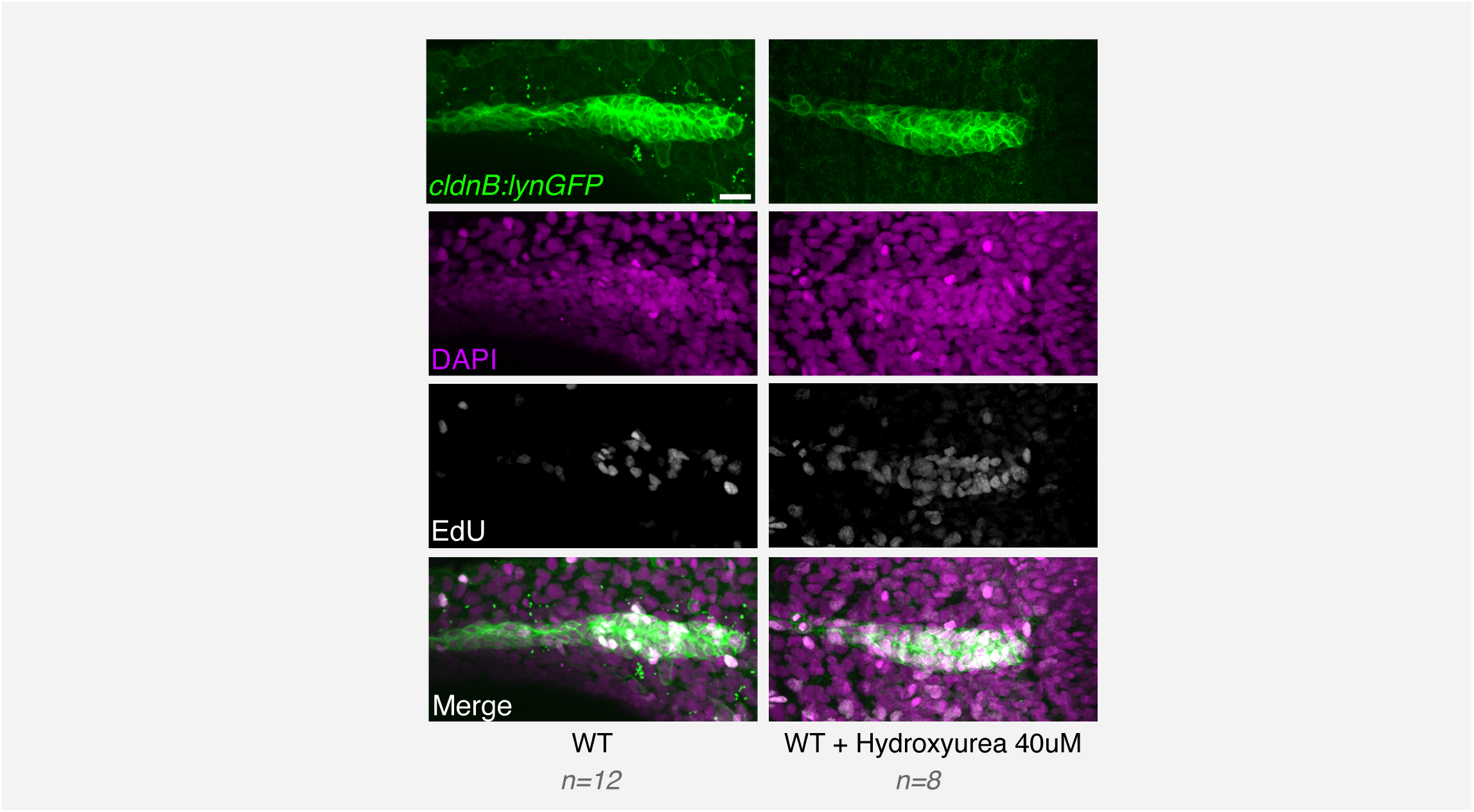
Hydroxyurea treatment results in an S-phase arrest. Z-projections of EdU-labeled pLL primordia in Control and 40mM Hydroxyurea treated embryos stained with *cldnb:lynGFP* and DAPI. Scale bars: 20 μm. Data are representative of at least two independent experiments.

**Supplementary Figure 13.**
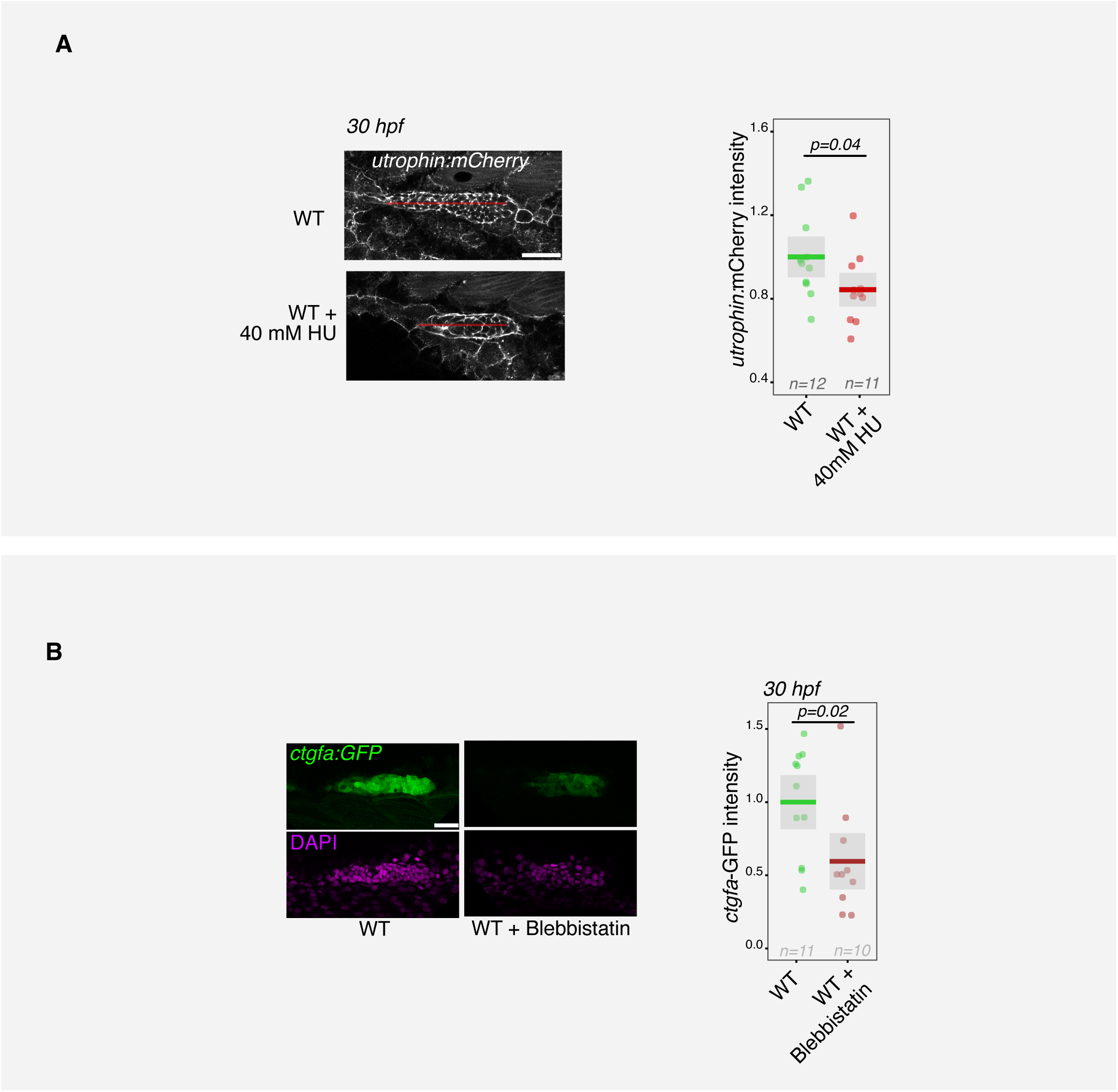
HU treatment induces diminished F-actin polymerization and actomyosin contractility controls primordium Yap/Taz signaling. **(A)** Z-stacks of pLL primordia from *act2b:Utrophin-mCherry* embryos treated with hydroxyurea. **(B)** Z-stacks of pLL primordia from *ctgfa:GFP* embryos treated with blebbistatin and stained with DAPI. Scale bars: 50 μm. Statistical analyses were performed using the unpaired Student’s t-test; *p-*values are indicated. Data are representative of at least two independent experiments.

**Video1**. Time-lapse movie of lateral line primordium depositing L2 and L3 neuromasts in WT (Top) and *sox2-/-* (Bottom) in the *Tg(cldnB:lynGFP)* background. Movies acquired with separate stage positions on microscope were stitched together for the purpose of representation. Embryos were imaged every 1 minute starting 25hpf. Every frame is a Maximum intensity projection.

**Video2**. Time-lapse movie of lateral line primordium depositing L2 and L3 neuromasts in WT (Top) and *yap-/-; taz+/-* (Bottom) in the *Tg(cldnB:lynGFP)* background. Movies acquired with separate stage positions on microscope were stitched together for the purpose of representation. Embryos were imaged every 1 minute starting 25hpf. Every frame is a Maximum intensity projection.

**Video3**. Time-lapse movie of lateral line primordium cell junction falling apart upon laser microdissection. Every frame is an individual z-stack. Laser microdissection was performed on 30hpf embryos. Scale Bar 5 μm.

